# An agent-based modeling approach for lung fibrosis in response to COVID-19

**DOI:** 10.1101/2022.10.03.510677

**Authors:** Mohammad Aminul Islam, Michael Getz, Paul Macklin, Ashlee N. Ford Versypt

**Affiliations:** Department of Chemical and Biological Engineering, University at Buffalo, The State University of New York, Buffalo, NY, 14260, USA; Department of Intelligent Systems Engineering, Indiana University, Bloomington, IN, 47408, USA; Department of Biomedical Engineering, University at Buffalo, The State University of New York, Buffalo, NY 14260, USA; Institute for Artificial Intelligence and Data Science, University at Buffalo, The State University of New York, Buffalo, NY 14260, USA

## Abstract

The severity of the COVID-19 pandemic has created an emerging need to investigate the long-term effects of infection on patients. Many individuals are at risk of suffering pulmonary fibrosis due to the pathogenesis of lung injury and impairment in the healing mechanism. Fibroblasts are the central mediators of extracellular matrix (ECM) deposition during tissue regeneration, regulated by anti-inflammatory cytokines including transforming growth factor beta (TGF-β). The TGF-β-dependent accumulation of fibroblasts at the damaged site and excess fibrillar collagen deposition lead to fibrosis. We developed an open-source, multiscale tissue simulator to investigate the role of TGF-β sources in the progression of lung fibrosis after SARS-CoV-2 exposure, intracellular viral replication, infection of epithelial cells, and host immune response. Using the model, we predicted the dynamics of fibroblasts, TGF-β, and collagen deposition for 15 days post-infection in virtual lung tissue. Our results showed variation in collagen area fractions between 2% and 40% depending on the spatial behavior of the sources (stationary or mobile), the rate of activation of TGF-β, and the duration of TGF-β sources. We identified M2 macrophages as primary contributors to higher collagen area fraction. Our simulation results also predicted fibrotic outcomes even with lower collagen area fraction when spatially-localized latent TGF-β sources were active for longer times. We validated our model by comparing simulated dynamics for TGF-β, collagen area fraction, and macrophage cell population with independent experimental data from mouse models. Our results showed that partial removal of TGF-β sources changed the fibrotic patterns; in the presence of persistent TGF-β sources, partial removal of TGF-β from the ECM significantly increased collagen area fraction due to maintenance of chemotactic gradients driving fibroblast movement. The computational findings are consistent with independent experimental and clinical observations of collagen area fractions and cell population dynamics not used in developing the model. These critical insights into the activity of TGF-β sources may find applications in the current clinical trials targeting TGF-β for the resolution of lung fibrosis.

**Author summary:** COVID-19 survivors are at risk of lung fibrosis as a long-term effect. Lung fibrosis is the excess deposition of tissue materials in the lung that hinder gas exchange and can collapse the whole organ. We identified TGF-β as a critical regulator of fibrosis. We built a model to investigate the mechanisms of TGF-β sources in the process of fibrosis. Our results showed spatial behavior of sources (stationary or mobile) and their activity (activation rate of TGF-β, longer activation of sources) could lead to lung fibrosis. Current clinical trials for fibrosis that target TGF-β need to consider TGF-β sources’ spatial properties and activity to develop better treatment strategies.

## Introduction

Severe acute respiratory syndrome coronavirus 2 (SARS-CoV-2) is responsible for the coronavirus disease 19 (COVID-19) pandemic and primarily causes pulmonary infection with symptoms and complications ranging from short-term pneumonia to long-term acute respiratory distress syndrome (ARDS) [1–3]. The long-term respiratory complications of COVID-19 survivors include diffuse alveolar damage (DAD), which is also a histological hallmark of ARDS [4]. DAD begins with the exudative phase when edema and inflammation occur, followed by the fibroproliferative phase with loose-organizing fibrosis. In the fibroproliferative phase, damaged tissues are partially replaced through fibroblast migration, proliferation, and collagen deposition. But impairments in the repair mechanisms—such as persistent infections [5], imbalances between profibrotic and antifibrotic mediators [6], and dysregulation in the functions of immune cells [7, 8] and fibroblasts [9, 10]—can lead to long-term fibrosis.

The severity of SARS-CoV-2 infection dictates the severity of pulmonary fibrosis. Zou et al. [11] reported that 100% of patients with severe and critical COVID-19 disease generate pulmonary fibrosis, wherein severe fibrosis is observed in most cases. Although 90-day follow-up results in their studies showed significant resolution in most patients, some patients had fibrotic areas after 90 days. In another study, Li et al. [2] reported that 59.6% of patients with confirmed COVID-19 cases had lung fibrosis after 90 days from onset. The overall lung fibrosis in hospitalized patients was 86.87%, out of which only 31.74% of patients had eventual resolution of fibrosis in follow-up visits. The six-month follow-up computed tomography scan of severe COVID-19 pneumonia survivors in the study of Han et al. [12] showed fibrotic changes in 35% of patients. They identified that susceptible groups have ARDS, higher heart rates, older age, longer hospital stays, and noninvasive mechanical ventilation. The pathological findings of COVID-19 pulmonary fibrosis identified alveolar epithelial cells (AECs) as the injury site. In contrast, other fibrotic lung diseases, e.g., idiopathic pulmonary fibrosis (IPF), can occur from epithelial and endothelial cell damage [3, 13]. Currently, the mechanistic basis for progressive and stable fibrosis resolution is not completely understood.

Macrophages play an essential role in different phases of DAD. Inflammatory macrophages (M1 macrophages) are activated during the exudative phase of tissue damage to provide host defense [14]. At the same time, M1 macrophages are responsible for inflammation and lung injury. In the fibroproliferative phase of DAD, M1 macrophages shift to alternatively activated macrophages (M2 macrophages) and start secreting anti-inflammatory cytokines (e.g., TGF-β), clearing apoptotic cells, and removing debris [15]. M2 macrophages play a crucial role in developing and resolving tissue damage in the fibroproliferative and fibrotic stages of DAD [16, 17].

Another critical contributor to fibrosis is the activation of latent TGF-β due to tissue damage. TGF-β is always secreted in a latent form that must be activated. Depending on the spatial behavior of the sources of TGF-β, they can be categorized into two groups: stationary sources (e.g., latent stores of TGF-β) and mobile sources (e.g., M2 macrophages). Excess TGF-β that is not activated within a short time of synthesis is stored in extracellular matrix (ECM) as latent deposits that are spatially restricted [18]. Latent TGF-β is stored at high concentrations in the ECM in an inactive state in most organs, including the lung [19]. Injury and death of epithelial cells due to external stimuli activate the latent TGF-β in the ECM at the damaged sites using αvβ6 integrins [9, 20]. Latent TGF-β can also be activated by reactive oxygen species, low pH, and proteolysis [21] and from cell surfaces [22]. Here, we refer to activation of TGF-β from the ECM stores without *de novo* synthesis as “stationary sources” of TGF-β.

Other sources of TGF-β in pulmonary fibrosis are M2 macrophages, metaplastic type II AECs, bronchial epithelium, eosinophils, fibroblasts, and myofibroblasts [21, 23]. In this work, *de novo* synthesis of latent TGF-β from anti-inflammatory M2 macrophages after clearance of dead epithelial cell debris [24] is assumed to be immediately activated; thus, this source of TGF-β is considered to be “mobile”. This is intended to capture the activation of latent TGF-β from cell surfaces including M2 macrophages [22, 25, 26].

Overexpression of TGF-β from any source and the persistent presence of any source can lead to fibrosis. Experimental observations in rat models showed the progression of pulmonary fibrosis with overexpression of TGF-β [27]. Also, quantification of TGF-β in the plasma of COVID-19 patients showed an increase in TGF-β concentration compared to healthy individuals [28, 29].

Activated TGF-β directs the functions of fibroblasts. Resident fibroblasts are located beneath epithelial cells or scattered in the ECM between the lung’s epithelial and endothelial layers [30]. The accumulation of TGF-β due to viral infection and epithelial cell injury recruits fibroblasts [31]. Small injuries can be repaired by the proliferation and differentiation of type 2 AECs to type 1 AECs. In contrast, the restoration process fails in severe injuries, leading to fibroblast activation and fibrosis [9]. Activated fibroblasts continually secrete ECM (e.g., collagen) to maintain the structural integrity of the lung [32]. TGF-β also regulates the collagen deposition rate of fibroblasts [33].

The current identification methods of lung fibrosis include histologic analysis of tissue samples and radiographic examination with computed tomography [34, 35]. Fibrotic pattern, collagen area fraction, and localization of cells can be observed from the histological analysis. Collagen area fractions evaluated from histopathological images are strongly correlated with histologic fibrosis scores and provide a reliable index to quantify fibrosis [36].

Existing mathematical and computational models can give insights into the fundamental processes that regulate fibrotic outcomes. Brown et al. [37] developed a simple agent-based model (ABM) considering the effects of the pro-inflammatory mediator TNF-α in lung tissue damage and the anti-inflammatory mediator TGF-β1 in fibroblast-mediated collagen deposition due to exposure to smoke particles. Their model captured the histological observation of self-resolving tissue damage, localized tissue damage with fibrosis, and extensive tissue damage with fibrotic states. The ABM of Warsinske et al. [38] explored the co-regulatory relationship between epithelial cells and fibroblasts through TGF-β1 during fibrosis. In another 3D ABM of lung fibrosis [39], the initial amount of damaged type 2 AECs activated the inactive TGF-β. The ABM of Ceresa et al. [40] focused on emphysema progression (excess degradation of collagen) in chronic obstructive pulmonary disease and considered the transition of macrophages between the M1 and M2 phenotypes, secretion of TGF-β from M2 macrophages, recruitment of fibroblasts by TGF-β, fibroblast-mediated collagen deposition, and collagen degradation by MMP9. The mathematical model of Hao et al. [41] used continuous partial differential equations for IPF and considered crucial cells and cytokines for the fibrosis process, including the transition of M1 macrophages to the M2 phenotype and TGF-β-mediated fibroblast proliferation. Their results showed how treatments that block the activity of cytokines (e.g., anti-TGF-β) could reduce the excess formation of ECM. While fibrosis in the lung and other organs such as the heart differ in etiology and outcome, many important similarities exist at the cellular and chemical levels between cardiac and pulmonary fibrosis [42]. Cardiac fibrosis is an area with rich experimental data and computational modeling studies. One study combined experiments and mathematical modeling to investigate the role of TGF-β in fibroblast-mediated collagen deposition during myocardial infarction (MI) [43]. Chowkwale et al. [44] also investigated inflammation-fibrosis coupling during MI considering the activation of latent TGF-β and TGF-β-dependent proliferation of fibroblasts. The coupled ABM and logic-based model of Rikard et al. [45] simulated similar increases in collagen area fractions to experimental observations during MI. Their model also considered the generation of latent TGF-β and fibroblast-mediated collagen deposition. In summary, TGF-β and fibroblasts are standard players in fibrosis progression in all the models described above, and major sources of TGF-β are M2 macrophages and latent stores of TGF-β. Although none of the models considered fibrosis due to viral infection, the process of fibrosis due to SARS-CoV-2 infection involves similar cells and chemical mediators.

We developed a multiscale lung tissue simulator that can be used to investigate the mechanisms of intracellular viral replication, infection of epithelial cells, host immune response, and tissue damage [46]. Here, we focused on the mechanisms of the fibrotic process in the lung tissue during SARS-CoV-2 infection. We conducted *in silico* experiments to determine the effects of the spatial behavior of TGF-β sources (stationary and mobile), the activation rate of TGF-β from sources, and the activation duration of TGF-β sources in the development and progression of fibrosis. We used collagen area fractions from histological analysis to compare the outcomes of our model simulations.

## Methods

### Overall model

In our earlier work, we developed a SARS-CoV-2 lung tissue simulator [46], which we refer to as the “overall model”. Here, we briefly recap the most salient features of this overall model before describing our new contribution: the “fibrosis model”.

The overall model structure is developed in an open-source multiscale ABM framework PhysiCell [47] . The model simulates a tissue section of 800 μm × 800 μm *×* 20 μm, representing a monolayer of stationary epithelial cells on an alveolar surface of lung tissue. Initially, 2793 epithelial cells, 50 resident macrophages, 28 dendritic cells (DCs), and 57 fibroblasts are present in the simulated tissue. These initializations are directly from the code used in the overall model manuscript [46]; here, we maintain the same cellular initializations. Cells are recruited to tissue through voxels that represent vasculature points; we randomly assign 8.8% voxels as vasculature points for arrivals of recruited cells because approximately this percentage of the tissue consists of the vasculature [48]. Cellular movement and diffusion of substrates occur along a 2D plane through or above the tissue, represented by a single layer of 3D voxels; this is consistent with the transport through thin alveolar tissue. We infect the simulated tissue by placing SARS-CoV-2 viral particles in the extracellular space using a uniform random distribution. The initial number of virions is selected based on the multiplicity of infection (MOI), defined as the ratio of initial virions to the number of epithelial cells. We used a default MOI = 0.1. Viral particles diffuse through tissue, bind with unoccupied ACE2 receptors on the epithelial cell surfaces, internalize via endocytosis, replicate through intracellular viral replication kinetics, and export back to the extracellular domain by exocytosis.

PhysiCell is coupled to BioFVM [49], which solves partial differential equations for chemicals considered to be continuous (as opposed to the discrete agents) with terms for diffusion, decay, uptake, and release of multiple substances simultaneously. The extracellular viral transport, uptake by adhering to cell surface ACE2, and virion export from the cell are modeled using BioFVM formulation with Neumann boundary conditions:

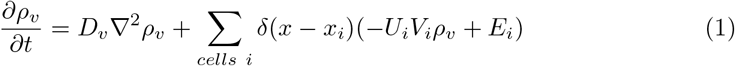

where *ρ*_*v*_ is the concentration or population density of the viruses, *D*_*v*_ is the diffusion coefficient, *U*_*i*_ is the virion uptake rate by the cell, *V*_*i*_ is the cell volume, and *E*_*i*_ is the virion export rate from the cell. We used *D*_*v*_ = 2.5 μm^2^min^-1^, which is within the range of influenza viral diffusion in the airway calculated in [50] (from 0.02 μm^2^min^-1^ to 190.8 μm^2^min^-1^). Moses et al. [51] also used a similar diffusion coefficient (1.88 μm^2^min^-1^) for SARS-CoV-2 lung infection.

Virions bind with unoccupied external ACE2 (R_EU_) using a continuum to discrete transition rule based on the binding flux:

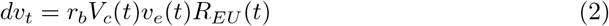

where *r*_*b*_ is the binding rate between virion and R_EU_, *V*_*c*_(*t*) is the volume of the cell, *v*_*e*_(*t*) is the density of the virions near a cell, and *R*_*EU*_ (*t*) is the number of R_EU_ at time *t*. The continuum to discrete transition rule allowed all nearby whole virions uptake when *dv*_*t*_ *>* 1. For fractional virion or for *dv*_*t*_ *<* 1, which resulted from the continuum rule in Eq 1, virion uptake depends on the probabilistic rule. The fractional virion uptake occurs when a random number drawn from a uniform distribution is greater than the fractional binding flux (*U*(0, 1) *> dv*_*t*_, where 0 *< dv*_*t*_ *<* 1).

The external virus-bound ACE2 receptors endocytose, release the virions, and recycle back to the cell surface. The intracellular released virions uncoated and synthesized viral RNA and proteins and assembled for exocytosis. The receptor kinetics and intracellular viral replication kinetics are modeled as a set of ordinary differential equations (ODEs), and corresponding parameters are selected from the experimental characterization of SARS-CoV entry into host cells [52]. The assembled virions also initiate cell death response, which is modeled as Hill function to relate the apoptosis rate of a cell to assembled virions. The infected cells secrete chemokine until the cell dies through lysis or CD8+ T cell-induced apoptosis. Details of the intracellular virus model for replication kinetics, viral response, and receptor trafficking are described in much greater detail in the overall model manuscript [53].

The resident and newly recruited macrophages move along the gradients of chemokine and debris, phagocytose dead cells, and secrete pro-inflammatory cytokines. We consider five states of macrophages: inactivated, M1 phenotype, M2 phenotype, hyperactive, and exhausted. Both M1 and M2 phenotypes can also exist in hyperactive and exhausted states. Neutrophils are recruited into the tissue by pro-inflammatory cytokines, move along the gradients of chemokine and debris, phagocytose dead cells, and uptake virions. The resident DCs chemotaxis along the gradients of chemokines, activated by infected cells and viruses, and a portion of the activated DCs egress out of the tissue to the lymph node to induce activation of virus-specific CD4+ and CD8+ T cells. The activated T cells are recruited in the tissue from the lymph node. CD8+ cells attempt to attach with infected cells based on PhysiCell’s mechanical interaction and cumulative attachment above a threshold time (25 min) causes apoptosis of the infected cells. The contact between CD4+ T cells and macrophages induces a hyperactive state in macrophages, enabling them to phagocytose live infected cells. The out-of-bound cells from the simulated domain are pushed back into the tissue section by changing the direction away from the edge.

Extracellular densities of chemokines and pro-inflammatory cytokines are modeled using standard BioFVM formulation with Neumann boundary conditions:

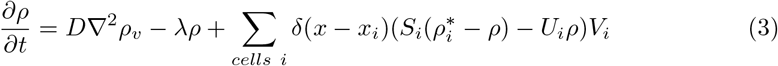

where *D* is the diffusion coefficient, *λ* is the decay rate, *S*_*i*_ is the secretion rate, *U*_*i*_ is the uptake rate, and 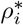 is the saturation density. For chemokines, pro-inflammatory cytokines, and debris, diffusion coefficients are assumed to be the same as the diffusion coefficient for monoclonal antibodies [54] and set at 555.56 μm^2^min^-1^; decay rates are assumed to be the same as the decay of IL-6 [55] and set at 1.02 *×* 10^−2^ min^-1^. Further details of parameters for immune cells, equations for chemotaxis and recruitment, rules for phagocytosis and CD8+ T cell-induced apoptosis, and proliferation, activation, and clearance of the T cells in the lymph nodes using ODEs are available elsewhere [46, 53, 56]. The equations for chemotaxis and cell recruitment are also defined in the following section as they are directly used in the fibrosis model. The overall model has 5 cell types and 76 biological hypotheses, listed in the Supplementary File, as adapted from the overall model manuscript [46]. The simulation update time for the microenvironment using BioFVM formulation is 0.01 min, cell mechanics is 0.1 min, cell processes is 6 min, and cell recruitment is 10 min. The total simulation time in the default case is 21,600 min (15 days), and the data output for saving every 60 min.

### Fibrosis model

We extended the overall model to include the mechanisms for fibrosis (Fig 1). We used the ABM platform PhysiCell to simulate the rules for phenotypic and chemical changes, including cell cycle, volume changes, death processes, cellular movement, and recruitment of the agents in Fig 1, in addition to the other processes included in the overall model. The rules for the following cell types in the fibrosis model (Fig 1)—epithelial cells, infected cells, CD8+ T cells, and M1 macrophages—are all used directly from the overall model [46].

**Fig 1.**
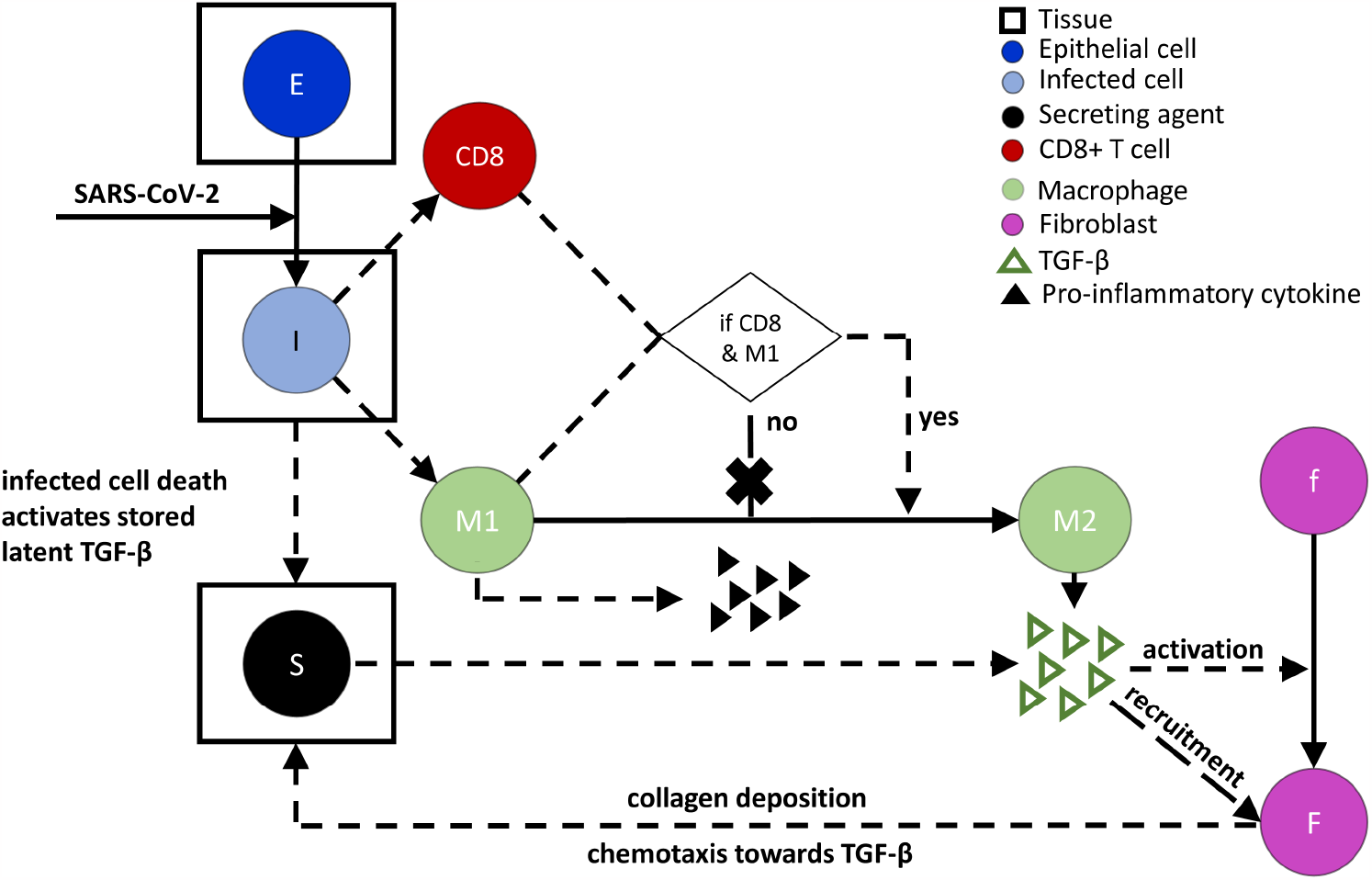
Cellular and chemical mediators in lung fibrosis during SARS-CoV-2 infection included in the fibrosis model. Secreting agents (S) mimic the activation of latent stores of TGF-β by acting as stationary sources of TGF-β for a certain period at the site and time of death of infected epithelial cells (I). Infected cells recruit M1 macrophages initially as an innate immune response and CD8+ T cells in the later phase of infection as an adaptive immune response. M1 macrophages stop secreting pro-inflammatory cytokines when located within close proximity to CD8+ T cells and instantaneously convert to M2 macrophages. M2 macrophages secrete and activate TGF-β and are considered mobile sources. Fibroblasts become activated and recruited by TGF-β, chemotax towards the gradient of TGF-β, and deposit collagen continuously. Cells with rectangles drawn around them represent stationary cells in tissue sections, while all other cells are considered to be mobile cells. Solid arrows denote the transformation of one cell species to another, dotted lines denote interactions that influence processes without the source being produced or consumed, and the X over an arrow indicates the process does not occur. Triangles represent secreted chemical factors that can diffuse and interact in the system. Cells with different color codes represent the corresponding cells in the simulation.

Several additional processes are included in the fibrosis model. These involve the secreting agents, M2 macrophages, inactive and activated fibroblasts, TGF-β, and collagen. The death of an infected epithelial cell (I in Fig 1) activates the latent TGF-β embedded in the tissue ECM. The activation of latent stores of TGF-β is represented by creating a stationary secreting agent (S in Fig 1) at the site and time of epithelial cell death. The secreting agents secrete TGF-β for a certain period as the amount of TGF-β stored in the ECM is limited; a death rate of secreting agents (*AT*) is used to represent terminating the activation of latent stores of TGF-β. Here, death rate *AT* is analogous to the extinction rate of the ECM-bound TGF-β sources and is inversely proportional to the mean duration of those sources.

Macrophages are the mobile sources of TGF-β in this fibrosis model. M1 macrophages are recruited in the initial phase, and CD8+ T cells are recruited in the later stage of infection. Usually, regulatory T cells (Tregs), a subset of CD4+ T cells, are involved in the phenotypic transformation of M1 to M2 macrophages [15]. Although we do not have Tregs in the current model, CD8+ T cells exhibit similar behavior by inhibiting the secretion of pro-inflammatory cytokines from M1 macrophages in the later phase of infection [46]. Since the fibroproliferative phase of DAD occurs in the later phase of the disease, a conditional rule is added in the fibrosis model to follow this pathology: if a CD8+ T cell and an M1 macrophage are within close proximity, then the M1 macrophage stops secreting pro-inflammatory cytokines, transforms to an M2 macrophage, and starts secreting TGF-β. The interaction distance for this transition is calculated using a multiple (*ϵ*) of the radii of the two interacting immune cells [46]. The transformation of the M1 to M2 phenotype is considered to occur instantaneously when the distance between an M1 macrophage and a CD8+ T cell is ≤ *ϵ* multiplied by the sum of the radii of the M1 macrophage and the CD8+ T cell. The cell radii are computed in PhysiCell from cell volumes over time as the cell volume changes with the progression of the cell cycle. The initial volume of macrophages is 4849 μm^3^ and that for CD8+ T cell is 478 μm^3^ with nuclear volumes of 10% of the cell volumes; these are the same values specified in the overall model [53].

We simulate extracellular concentrations of TGF-β (*T*_*β*_) using the standard BioFVM formulation with Neumann boundary conditions:

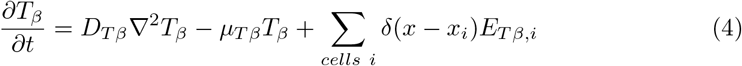

where *t* is time, *D*_*T β*_ is the diffusion coefficient of TGF-β, *μ*_*T β*_ is the net decay and bulk removal rate, *δ*(*x*) is the discrete Dirac delta function, *x* is the center of the voxel, *x*_*i*_ is the position of the center of discrete cell *i*, and *E*_*T β,i*_ is the net export rate of TGF-β from cell *i*. Net export rate *E*_*T β,i*_ denotes activation rate of latent stores of TGF-β at the damaged site, which we term damaged-site secretion (*DS*), or TGF-β secretion and activation rate from macrophages, which we term macrophage secretion (*MS*).

Initial fibroblasts remain in an inactive state. The experiments of mice model of MI showed a homeostatic value of fibroblasts without external perturbation or injury [57]. To maintain the homeostatic population of fibroblasts, we assume that the initial inactive fibroblasts move randomly throughout the domain and do not undergo apoptosis (i.e., *μ*_*F*_ = 0). TGF-β activates the resident inactive fibroblasts. We assume that inactive fibroblasts become active fibroblasts in the presence of TGF-β (*T*_*β*_ *>* 0) and switch back to the inactive state in the absence of TGF-β (*T*_*β*_ = 0). The active fibroblasts and inactive fibroblasts from the active state undergo apoptosis naturally at a rate of *μ*_*F*_ and become dead cells. Both inactive and active fibroblasts are subject to the cycle cell model in PhysiCell where their cell volumes are over updated time as the cell cycle progresses. The initial volume of a fibroblast (*F*_*V*_) and the corresponding nucleus (*F*_*V N*_) are specified.

The activated fibroblasts chemotax up the gradient of TGF-β. The chemotaxis velocity of a fibroblast 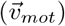 depends on the TGF-β concentration in its neighboring regions, fibroblast speed (*s*_*mot*_), migration bias (*b*), and time for persistence in a specific trajectory (∆*t*_*mot*_) [46]:

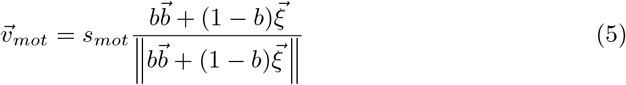

where *s*_*mot*_ is the speed of chemotaxis, 0 ≤ *b* ≤ 1 is the level of migration bias (i.e., *b* = 0 represents no influence of chemotaxis and only random cell migration),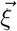 is a random unit vector direction, and 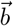 is the migration bias direction. Fibroblasts also persist on their given trajectory for ∆*t*_*mot*_ before a new trajectory is computed. For fibroblasts,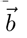 is estimated from the gradient of TGF-β concentrations in the neighboring regions [46]:

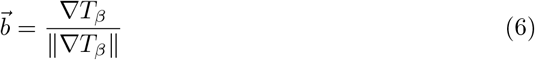

Chemotaxis of other immune cells along the gradients of pro-inflammatory cytokines, chemokines, and debris in the overall model follows this formulation [46].

New fibroblasts are recruited into the tissue by TGF-β. The number of fibroblasts recruited to the tissue (*N*_*r*_) is determined by integrating the recruitment signal over the whole domain:

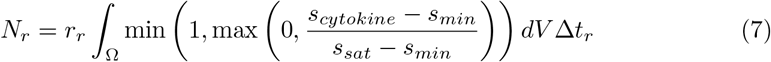

where Ω is the computational domain, *r*_*r*_ is the recruitment rate (per volume), *s*_*min*_ is the minimum recruitment signal, *s*_*sat*_ is the saturating or maximum signal, and *s*_*cytokine*_ is the recruitment signal that depends on the cytokine concentration. The volume of each grid voxel is *dV* = 20 *×* 20 *×* 20 μm^3^, and the time interval for recruitment of fibroblasts is ∆*t*_*r*_ = 10 min, the same as in the overall model [46]. Eq 7 gives the number of fibroblasts recruited between *t* and *t* + ∆*t*_*r*_. Here, *s*_*cytokine*_ = *F*_*g*_(*T*_*β*_) for the recruitment signal of fibroblasts via the cytokine TGF-β. Recruitment of any immune cell by a specific cytokine in the overall model follows this formulation; proinflammatory cytokine-dependent recruitment of neutrophils and macrophages is already included in the overall model [46].

The function for TGF-β-dependent recruitment of fibroblasts (*F*_*g*_(*T*_*β*_))—particularly its specific parameters—comes from previously published works [40, 43]. The parameters are estimated based on experimental data and illustrated in Fig S1A. We selected a threshold value to set the upper range of TGF-β concentration beyond which *F*_*g*_(*T*_*β*_) is assumed to be constant because of the polynomial nature of the recruitment signal.

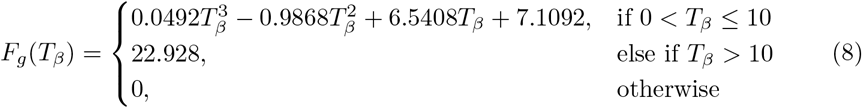

The activated fibroblasts deposit collagen continuously. The newly deposited collagen is assumed to not diffuse or degrade. TGF-β-dependent collagen deposition from fibroblasts is described by

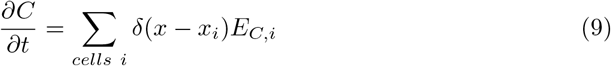

and

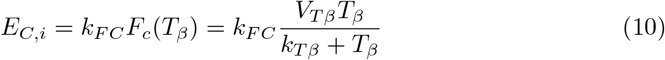

where *C* is the concentration of collagen, *E*_*C,i*_ is the net export rate of collagen from fibroblast cell *i*, and *k*_*F C*_ is the collagen production rate by fibroblasts. *F*_*c*_(*T*_*β*_) defines the TGF-β dependence of fibroblast-mediated collagen deposition (Fig S1B), which is assumed to follow a Michaelis-Menten form, where *V*_*T β*_ and *k*_*T β*_ are the Michaelis-Menten limiting rate and half saturation parameters, respectively.

### Parameter estimation

Experimental data from the literature used in this section were extracted from graphs in the original sources (referenced in the text) using the open-source Java program Plot Digitizer [58]. The initial values of the species in the uninfected condition are listed in Table 1. The estimated and calculated parameters are listed in Table 2. Several parameters were taken from the literature without new parameter estimation: *AT, ϵ, D*_*T β*_, *μ*_*T β*_, *μ*_*F*_, *F*_*V*_, *F*_*V N*_, *V*_*F*_, *s*_*mot*_, *b*, ∆*t*_*mot*_, and *r*_*r*_. Their values and sources are also listed in Table 2. *AT* was also varied in one of the *in silico* experiments.

**Table 1.**
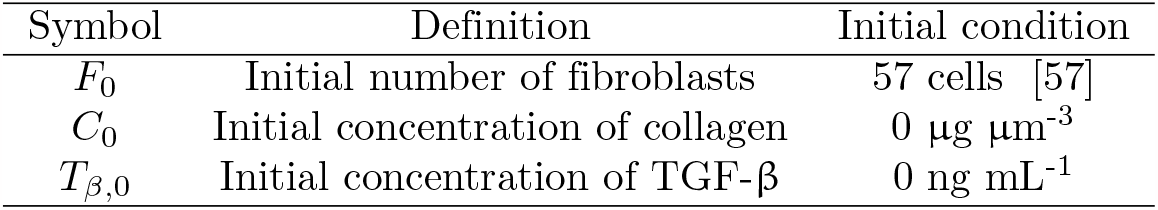
Initial conditions for model variables.

**Table 2.** Parameters for the fibrosis model.

| Symbol | Definition | Value | Unit | Source |
| --- | --- | --- | --- | --- |
| $AT$ | Activation time of TGF-β, via secreting agent death rate | $1 \times 10^{-3}$ | min$^{-1}$ | estimated from [43] |
| $\epsilon$ | Multiple of interaction distance for immune cells | 1.75 | Unitless | [46] |
| $D_{T\beta}$ | TGF-β diffusion coefficient | 3000 | $\mu\text{m}^2 \text{ min}^{-1}$ | [41] |
| $\mu_{T\beta}$ | TGF-β degradation rate | 15 | day$^{-1}$ | [40] |
| $DS$ | Damaged-site secretion rate of TGF-β | $5 \times 10^{-10} - 2 \times 10^{-9}$ | ng min$^{-1}$ | estimated |
| $MS$ | Macrophage secretion rate of TGF-β | $5 \times 10^{-10} - 5 \times 10^{-9}$ | ng min$^{-1}$ | estimated |
| $\mu_F$ | Active fibroblast apoptosis rate | $8.30 \times 10^{-5}$ | min$^{-1}$ | [40] |
| $F_V$ | Fibroblast cell volume | 2500 | $\mu\text{m}^3$ | [59] |
| $F_{VN}$ | Fibroblast nucleus volume | 500 | $\mu\text{m}^3$ | [60] |
| $V_F$ | Velocity of fibroblasts | 1 | $\mu\text{m min}^{-1}$ | [61] |
| $s_{mot}$ | Speed of fibroblasts | 1 | $\mu\text{m min}^{-1}$ | [61] |
| $b$ | Fibroblast migration bias | 0.7 | Unitless | [46] |
| $\Delta t_{mot}$ | Fibroblast directional persistence | 5 | min | [53] |
| $r_r$ | Recruitment rate of fibroblasts | $4 \times 10^{-9}$ | cells $\mu\text{m}^{-3} \text{ min}^{-1}$ | [46] |
| $s_{min}$ | Minimum fibroblast recruitment signal of TGF-β | 7.1092 | ng mL$^{-1}$ | calculated by Eq 8 |
| $s_{sat}$ | Maximum/saturating fibroblast recruitment signal of TGF-β | 22.928 | ng mL$^{-1}$ | calculated by Eq 8 |
| $k_{FC}$ | Fibroblast collagen production rate | $2.52 \times 10^{-7}$ | $\mu\text{g min}^{-1}$ | estimated from [41] |
| $V_{T\beta}$ | Rate parameter for TGF-β-dependent collagen deposition | $9.42 \times 10^{-1}$ | Unitless | estimated from [62, 63] data |
| $k_{T\beta}$ | Saturation parameter for TGF-β-dependent collagen deposition | $1.74 \times 10^{-1}$ | Unitless | estimated from [62, 63] data |
| $U_F$ | Uptake rate of TGF-β by each fibroblast | $1 \times 10^{-1}$ | min$^{-1}$ | estimated |

The initial number of fibroblasts (*F*_0_) was selected based on *in vivo* experiments of mice models of MI [57]. We extracted the fibroblast population per surface area from the experimental control condition and multiplied it by the surface area of the computational domain (800 *×* 800 μm^2^) to evaluate the initial number of inactive resident fibroblasts in the simulated tissue. No fibroblasts are activated at *t* = 0. We considered the concentrations of TGF-β (*T*_*β*_) and collagen deposited (*C*) to be changed from homeostatic values and set them to 0 as initial values (Table 1).

We evaluated the signal of TGF-β to recruit new fibroblasts (*s*_*cytokine*_) in Eq 7 using TGF-β-dependent recruitment of fibroblasts rate from Eq 8 (Fig S1A), i.e., *s*_*cytokine*_ = *F*_*g*_(*T*_*β*_). The minimum and maximum (i.e., saturating) recruitment signals (*s*_*min*_ and *s*_*sat*_, respectively) were calculated using Eq 8 for the lower (0 ng/mL) and upper (10 ng/mL) ranges of TGF-β, respectively.

The fibroblast collagen production rate (*k*_*F C*_) was estimated from Hao et al. [41] (details in the Supplementary File). *F*_*c*_(*T*_*β*_) is the TGF-β dependent collagen deposition rate. We normalized published experimental data [62, 63] and fitted the data to a Michaelis-Menten relationship to estimate the parameters *V*_*T β*_ and *k*_*T β*_ for the *F*_*c*_(*T*_*β*_) formulation in Eq 10 (Fig S1B).

The TGF-β-dependent functions (Fig S1) for recruitment rate of fibroblasts (*F*_*g*_(*T*_*β*_)) and collagen deposition rate from fibroblasts (*F*_*c*_(*T*_*β*_)) were defined for TGF-β ranges of 0–10 ng/mL from experiments [62–64] and mathematical models [40, 43]. We calibrated the fibrosis model by adjusting the free parameters *DS* and *MS* to keep TGF-β concentration within the defined range (Fig S2). We iteratively changed *DS* while *MS* = 0 (turning off secretion from macrophages) and changed *MS* while *DS* = 0 (turning off secretion/activation from damaged sites) to estimate the concentration range of TGF-β (Fig S2); our goal was to keep TGF-β concentration within the range of experimental observations (0–10 ng/mL) where the TGF-β-dependent functions for fibroblasts were defined. We selected ranges of *DS* from 5 *×* 10^−10^ to 2 *×* 10^−9^ ng/min and *MS* from 5 *×* 10^−10^ to 5 *×* 10^−9^ ng/min (Table 2).

## Statistical analysis

The statistical significance is assessed by the Mann-Whitney-Wilcoxon test two-sided with Bonferroni correction using the statannot package in Python [65]. *p* values are represented by: not significant (ns) *p >* 0.05; \**p* ≤0.05; \*\**p* ≤0.01; \*\*\**p* ≤0.001; \*\*\*\**p* ≤0.0001.

Each *in silico* experiment is replicated 15 times to capture the variability. The variability arises from the initial random placement of virions, random assignment of vasculature points for immune cell entry, uptake of virion by epithelial cells, distribution of cytokines, chemokines, and debris in the microenvironment and number of recruited immune cells based on that, and migration bias in cell velocity. We used mean, 5th percentile, and 95th percentile to show the variations in replications.

## Computational expense

All the simulations are performed in a Dell Precision 3640 tower workstation: Intel Core Processor i9-10900K (10 core, 20 MB cache, base 3.7 GHz, up to 5.3GHz, and 32GB RAM) using hyperthreading, for 6 total execution threads. For a single run with 21600 minutes of simulation, the total wall clock time was around 18 minutes.

## Code availability

The repository for version 5.0 of the overall model is available at https://github.com/pc4covid19/pc4covid19/releases/tag/v5.0 [66]. We have provided the code and analysis files for the fibrosis model in a repository at https://github.com/ashleefv/covid19fibrosis [67].

## Results and Discussion

We used the fibrosis model to predict the effects of stationary and mobile sources of TGF-β in fibroblast-mediated collagen deposition in the lung tissue in response to SARS-CoV-2 infection. The overall model produced predictions for the epithelial and infected cells, virions, and immune cells. The dynamics of these species are inputs to the fibrosis model. The fibrosis model output does not feedback to interact with any of the species of the overall model directly. A subset of the species not explicitly considered in the fibrosis model is shown in Fig S3. The fibrosis model was developed considering TGF-β activation from stationary agents to simulate the dynamics of TGF-β activation from latent stores in the ECM at the epithelial damage site. Also, TGF-β secretion from mobile sources simulated the contribution from M2 macrophages. For our default case, we considered equal TGF-β activation and secretion rates from both sources (case DM, Table 3) within the physiological range of TGF-β. We analyzed the spatial behavior of both sources and compared them with independent experimental and clinical observations. We also conducted a series of *in silico* experiments by turning on/off TGF-β activation and secretion rates from stationary and mobile sources, increasing/decreasing the rates from sources in combination, increasing the duration of the sources, and including additional rules for removal of TGF-β by chemotaxing fibroblasts. We validated our model with independent experimental data from mouse models for the dynamics of TGF-β, collagen area fraction, and macrophage cell population. We listed all the *in silico* experiments performed for this study in Table 3.

**Table 3.**
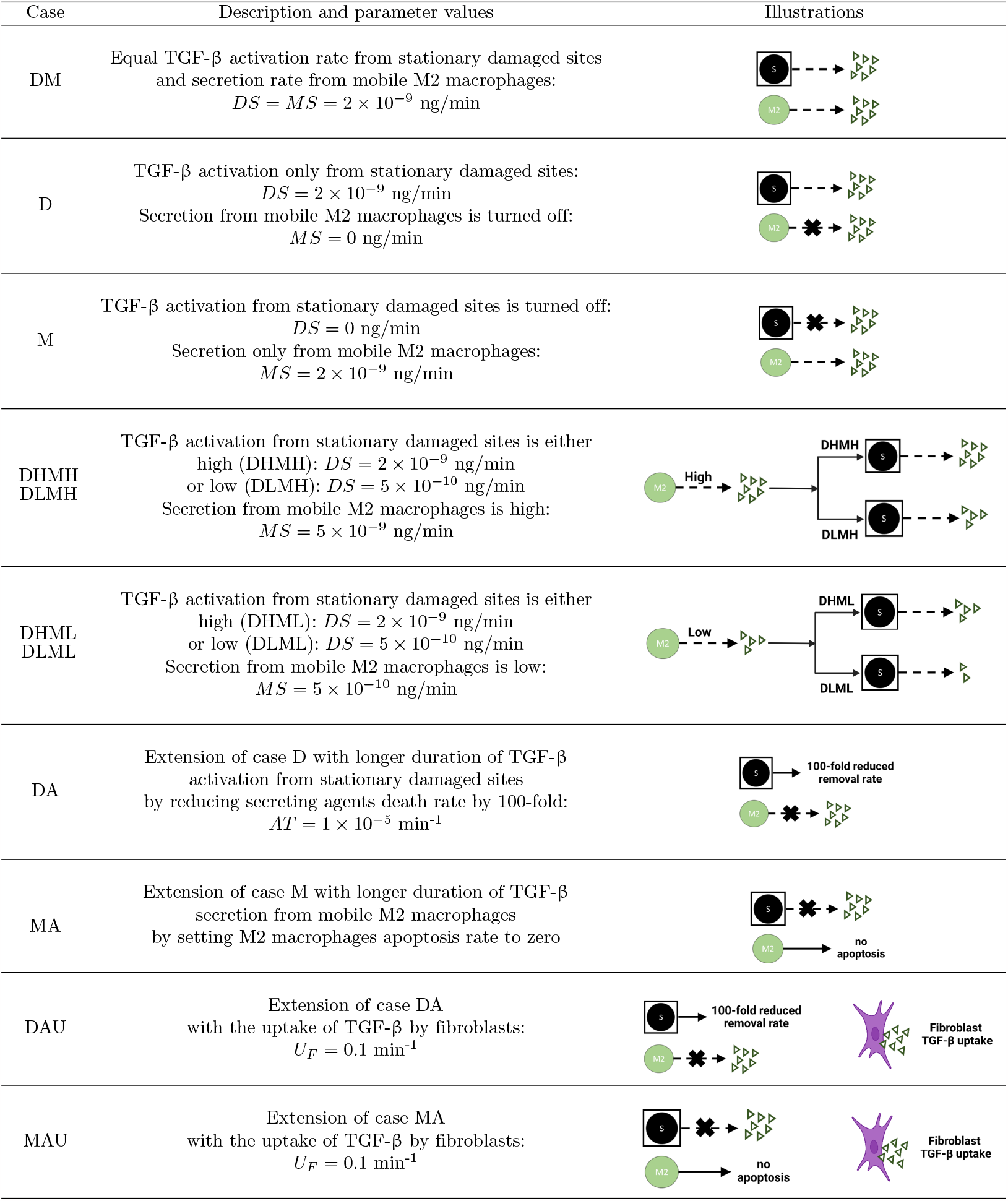
Details for *in silico* experiment cases. Note that the values of parameters from Table 2 were used unless otherwise specified below.

| Case | Description and parameter values | Illustrations |
| --- | --- | --- |
| DM | Equal TGF-β activation rate from stationary damaged sites and secretion rate from mobile M2 macrophages:<br>$DS = MS = 2 \times 10^{-9}$ ng/min | |
| D | TGF-β activation only from stationary damaged sites:<br>$DS = 2 \times 10^{-9}$ ng/min<br>Secretion from mobile M2 macrophages is turned off:<br>$MS = 0$ ng/min | |
| M | TGF-β activation from stationary damaged sites is turned off:<br>$DS = 0$ ng/min<br>Secretion only from mobile M2 macrophages:<br>$MS = 2 \times 10^{-9}$ ng/min | |
| DHMH<br>DLMH | TGF-β activation from stationary damaged sites is either high (DHMH): $DS = 2 \times 10^{-9}$ ng/min or low (DLMH): $DS = 5 \times 10^{-10}$ ng/min<br>Secretion from mobile M2 macrophages is high:<br>$MS = 5 \times 10^{-9}$ ng/min | |
| DHML<br>DLML | TGF-β activation from stationary damaged sites is either high (DHML): $DS = 2 \times 10^{-9}$ ng/min or low (DLML): $DS = 5 \times 10^{-10}$ ng/min<br>Secretion from mobile M2 macrophages is low:<br>$MS = 5 \times 10^{-10}$ ng/min | |
| DA | Extension of case D with longer duration of TGF-β activation from stationary damaged sites by reducing secreting agents death rate by 100-fold:<br>$AT = 1 \times 10^{-5}$ min$^{-1}$ | |
| MA | Extension of case M with longer duration of TGF-β secretion from mobile M2 macrophages by setting M2 macrophages apoptosis rate to zero | |
| DAU | Extension of case DA with the uptake of TGF-β by fibroblasts:<br>$U_F = 0.1$ min$^{-1}$ | |
| MAU | Extension of case MA with the uptake of TGF-β by fibroblasts:<br>$U_F = 0.1$ min$^{-1}$ | |

### Spatial behaviors of TGF-β sources affect collagen dynamics

For the spatial behaviors, we considered the cases of DM, D, and M (Table 3). Figs S4–S6 show the population spatial distributions over time for each agent type for these three cases. The stationary sources caused localized collagen deposition, whereas the distribution of collagen was more dispersed for mobile sources (Fig 2). We also observed two distinct peaks in TGF-β dynamics depending on the sources. Initially, the death of infected epithelial cells created secreting agents, which were stationary sources that activated latent TGF-β stored in the ECM at the damage sites. The number of these secreting agents fixed at damaged sites had a maximum around 5 days post-infection. The population reduced after that as the removal of the virions from the system occurred around 6 days (Fig S3A). As a result, we observed the first peak in TGF-β concentration around 5 days. The constant rate of activation and diffusion of TGF-β from stationary sources created localized fields with higher TGF-β concentrations. Active fibroblasts crowded at the sites of higher concentration of TGF-β and deposited collagen to cause localized distribution. In this model, M2 macrophages appear after the arrival of CD8+ T cells in the later phase of infection as we have encoded the rule that the interactions between M1 macrophages and CD8+ T cells transform M1 macrophages into the M2 phenotype, which acts as mobile sources of TGF-β. We observed the maximum number of macrophages around 9 days, which caused the second peak in the TGF-β profile. In the later phase of infection, damage was already present in the tissue when M2 macrophages arrived. M2 macrophages moved along the edge of damaged sites, phagocytosed dead and infected cells, cleared viral particles, and simultaneously secreted TGF-β. The movement of M2 macrophages along the edge of damaged sites caused higher concentrations of TGF-β at those sites. As a result, fibroblasts chemotaxed to those sites and deposited collagen. The deposited collagen was more dispersed with a higher collagen area fraction in the cases (DM and M) with mobile sources of TGF-β secretion compared to those (case D) with only stationary sources of TGF-β activation.

**Fig 2.**
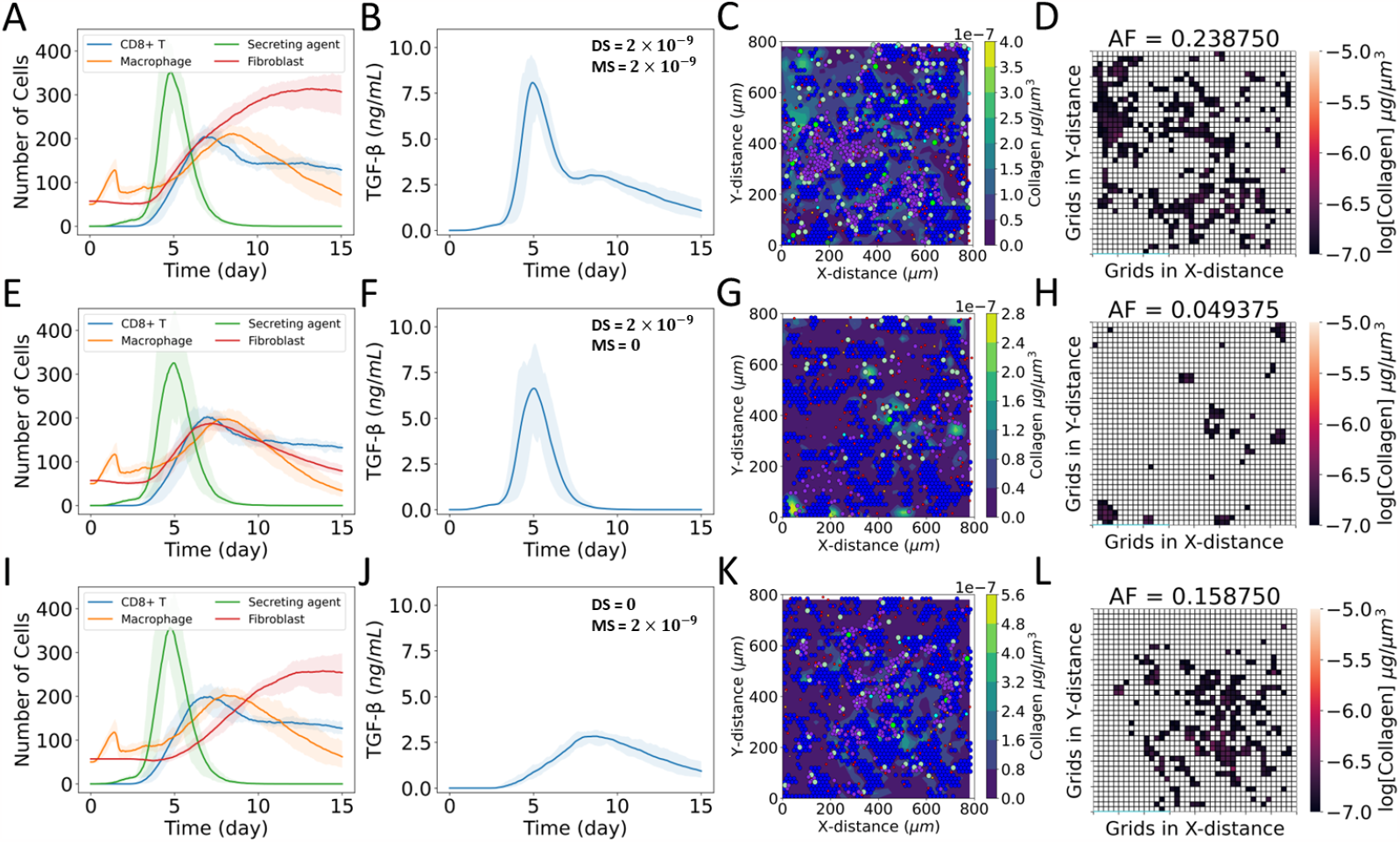
Effects of stationary and mobile sources of TGF-β on the dynamics of collagen deposition. (A–D) Case DM: equal TGF-β activation rate from stationary damaged sites and secretion rate from mobile macrophages, (E–H) Case D: TGF-β activation from stationary damaged sites only, and secretion from mobile macrophages is turned off, (I–L) Case M: TGF-β secretion from mobile macrophages only, and activation rate from stationary damaged sites is turned off. (A, E, I) Dynamics of populations of CD8+ T cells, macrophages, latent TGF-β activation sites (secreting agents), and fibroblasts; (B, F, J) dynamics of TGF-β concentration spatially averaged over the domain; (C, G, K) collagen deposition at damaged sites of tissue at day 15; (D, H, L) heat map showing collagen area fraction (AF) above the threshold value of 1 *×*10^−7^ μg μm^-3^. The solid curves represent the mean of predictions, and shaded areas represent the predictions between the 5th and 95th percentile of 15 replications of the agent-based model. The cases are defined in Table 3.

### Case DM: equal TGF-β activation rate from stationary damaged sites and secretion rate from mobile macrophages

In case DM (Fig 2A and Fig S4), we observed localization of fibroblasts around 2 days (Fig S4B,I) due to the slow creation of new stationary sources before the inflammatory response infiltrated the tissue. Around 5 days (Fig S4C,J), the sudden increase of stationary sources and the appearance of mobile sources (Fig 2A) disrupted that localization. Fibroblasts started to move along the edge of damaged sites as the mobile sources moved along the edge to clear infected and dead cells that were more prominent from 9 days to 15 days (Fig S4F,G,M,N). Fibroblasts had partial localization between 5 days and 9 days (Fig S4C–F,J–M). However, the spatial positions of localized fibroblasts changed with the disappearance of stationary sources and the appearance of mobile sources.

### Case D: TGF-β activation from stationary damaged sites only

In case D (Fig 2E and Fig S5), as in case DM, the slow creation of new stationary sources resulted in fibroblast localization at damaged sites around 2 days (Fig S5B,I). The rapid creation of new sources disrupted the localization of fibroblasts around 5 days (Fig S5C,J). Localization of fibroblasts was again observed between 5 days and 9 days (Fig S5C–F,J–M) followed by delocalization of fibroblasts (Fig S5F,G,M,N) as stationary sources stopped secreting TGF-β (Fig 2F). The absence of TGF-β deactivated fibroblasts, halting fibroblast-mediated collagen deposition after 9 days (Fig S5F,G).

### Case M: TGF-β secretion from mobile macrophages only

In case M (Fig 2I and Fig S6), we did not observe the localization of fibroblasts around 2 days (Fig S6B,I) when the TGF-β activation rate from damaged sites was turned off. Around 5 days (Fig S6C,J), fibroblasts were observed to move along the edge of damaged areas with the appearance of mobile sources of TGF-β (Fig 2I), which were more prominent from 9 days to 15 days. Overall, the TGF-β concentration for case M (Fig 2J) was less than that for cases DM (Fig 2B) and D (Fig 2F).

### Comparison between simulated and clinical studies of collagen area fraction

In summary for cases DM, D, and M, we observed changes in fibrotic patterns depending on the partial creation and deletion of sources, mobility of the sources, and days post infection. The combination of stationary and mobile sources of TGF-β in different phases of infection created higher TGF-β concentration zones at more sites than a single source type, resulting in higher collagen area fractions (Fig 2D,H,L). The distribution of collagen area fractions for cases DM, D, and M are shown in the violin plots in Fig 3. The collagen area fraction was lower in case D, intermediate in case M, and higher in case DM; Fig S7 shows the dynamics of the spatially averaged collagen concentrations for cases DM, D, and M. We observed exponential increases of spatially averaged collagen concentration in cases DM and M with a steeper slope in case DM; case D reached a plateau.

**Fig 3.**
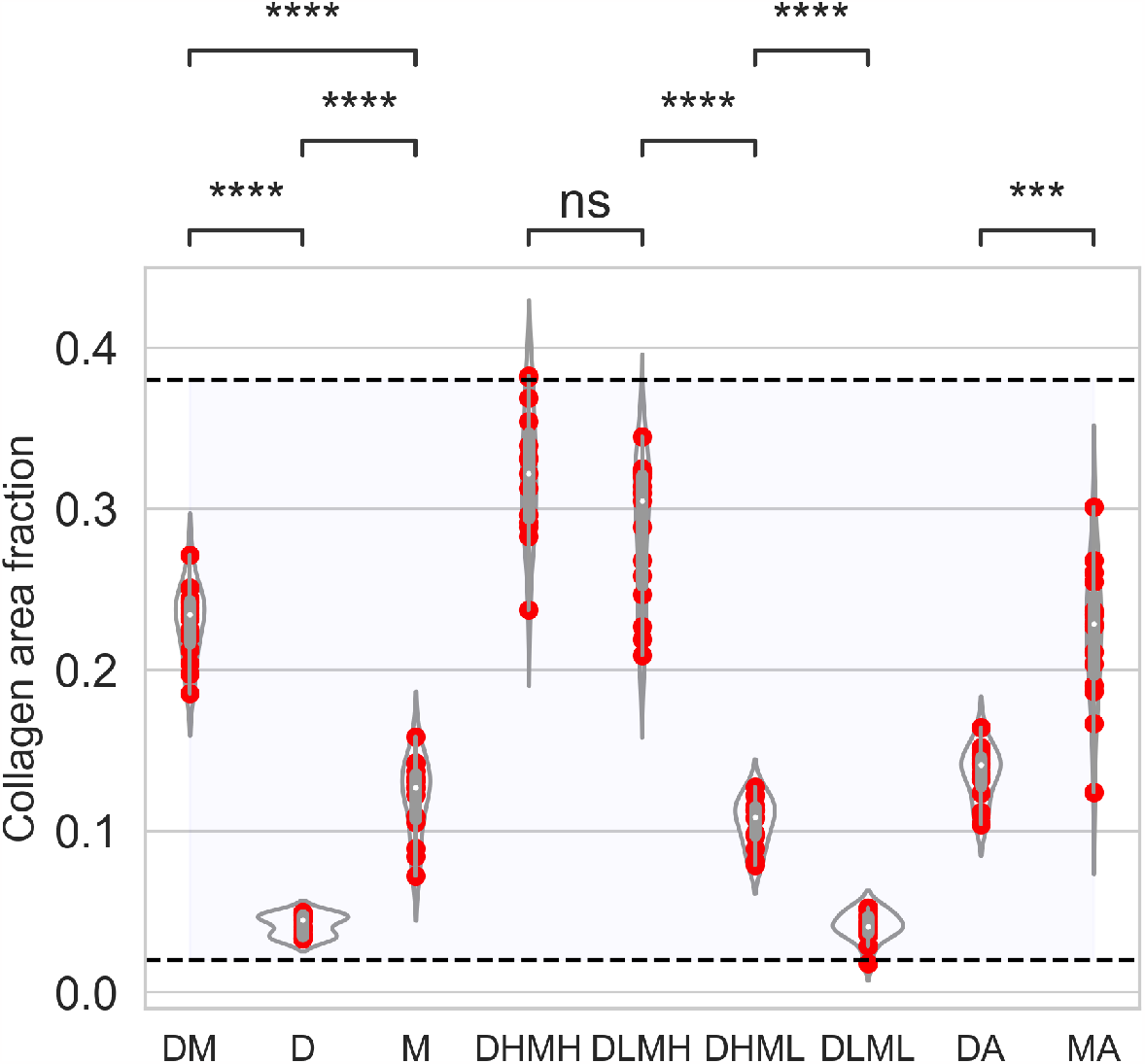
Quantification of collagen area fraction for different cases. Violin plots represent distribution, red solid circles represent data for 15 replications of each of the cases, and box plots show medians and quartiles. Case DM: equal TGF-β production rate from both damaged sites and macrophages; case D: TGF-*β* activation from damaged sites only and secretion from macrophages are turned off; case M: TGF-*β* secretion from macrophages only and activation rate from damaged sites are turned off; case DHMH: TGF-*β* activation rate from damaged sites is high and TGF-*β* secretion rate from macrophages is high; case DLMH: TGF-*β* activation rate from damaged sites is low and TGF-*β* secretion rate from macrophages is high; case DHML: TGF-*β* activation rate from damaged sites is high and TGF-*β* secretion rate from macrophages is low; case DLML: TGF-*β* activation rate from damaged sites is low and TGF-*β* secretion rate from macrophages is low; case DA: longer activation length of latent TGF-*β* in case D; case MA: longer activation length of M2 macrophages in case M. The horizontal dashed lines and shading indicate the ranges of collagen extension reported in Ball et al. [34]. The cases listed along the x-axis are defined in Table 3. *p* values are represented by: ns *p >* 0.05; \**p* ≤ 0.05; \*\**p* ≤ 0.01; \*\*\**p* ≤ 0.001; \*\*\*\**p* ≤ 0.0001.

Literature values for measurements performed on postmortem transbronchial cryobiopsy samples of COVID-19 pneumonia diseased tissue were reported with 2%–38% collagen extension considering all the lobes of the lung [34]. Analogously to our simulation domain, the percentages of collagen in the samples were determined by restricting the samples to the alveolar tissue while excluding airway walls, vessels, and airspaces. Our simulated collagen area fractions (Fig 3) are in similar ranges (2%–40%) compared to the experimentally observed percentage of collagen extensions at the endpoint outcomes. Our simulated collagen area fractions are also in a similar range to the reported increase of collagen volume fraction in human lung with different stages of IPF [68] and predicted collagen area fractions over time in the multiscale model and experiments of MI [45]. Although the mechanisms for fibrosis may differ in these analyses, our findings suggest that the activity of TGF-β sources is an important aspect, which alone can generate fibrotic areas similar to experimental observations.

### Higher TGF-β secretion rate from M2 macrophages leads to higher collagen area fraction

We explored the effects of varying the TGF-β activation rate from stationary damaged sites and the secretion rate from mobile macrophages at different levels in combination: cases DHMH, DLML, DHML, and DLMH (Table 3). Fig 4 shows the impacts of these variations, particularly on fibroblasts, TGF-β dynamics spatially averaged over the domain, and the TGF-β impact on the distribution of collagen deposition. Figs S8–S11 show the population spatial distributions over time for each agent type for these four cases.

**Fig 4.**
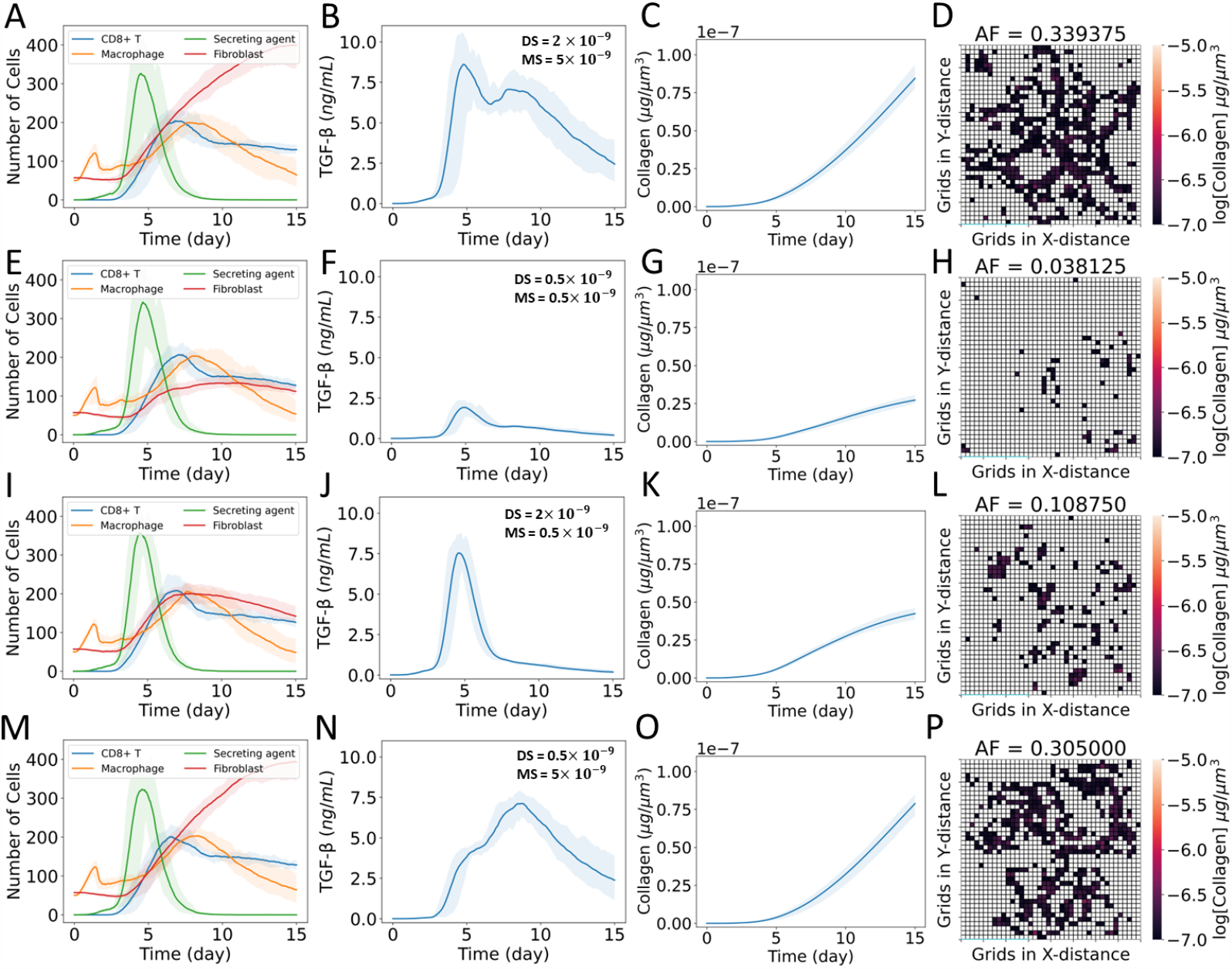
Effects of varying the TGF-β activation rate from stationary damaged sites *DS* and the secretion rate from mobile macrophages *MS* at different levels in combination. H and L denote higher and lower values, respectively, of *DS* when they come after D and of *MS* when they come after M in the case names. (A–D) Case DHMH, (E–H) case DLML, (I–L) case DHML, and (M-P) case DLMH. (A, E, I, M) Dynamics of populations of CD8+ T cells, macrophages, latent TGF-β activation sites (secreting agents), and fibroblasts; (B, F, J, N) dynamics of TGF-β concentration spatially averaged over the domain; (C, G, K, O) dynamics of collagen concentration spatially averaged over the domain; and (D, H, L, P) heat map showing collagen area fraction (*AF*) above the threshold value of 1 *×* 10^−7^ μg μm^-3^. The solid curves represent the mean of predictions, and shaded areas represent the predictions between the 5th and 95th percentile of 15 replications of the agent-based model. The cases are defined in Table 3.

### Cases DHMH and DLMH: TGF-β secretion from mobile macrophages is high and activation from stationary damaged sites is high/low

The agent dynamics showed high recruitment of fibroblasts for case DHMH (Fig 4A and Fig S8) and case DLMH (Fig 4M and Fig S11). We observed the highest collagen area fraction in case DHMH followed by case DLMH (Fig 3 and Fig 4D,P). In both cases, the TGF-β secretion rate from M2 macrophages was high. The spatially averaged TGF-β concentration remained high for the majority of the time period in both cases (Fig 4B,N) leading to very similar spatially averaged collagen concentrations (Fig 4C,O). Thus, mobile sources of TGF-β played a crucial role in higher collagen area fraction.

### Cases DHML and DLML: TGF-β secretion from mobile macrophages is low and activation from stationary damaged sites is high/low

In both cases DHML and DLML when the TGF-β secretion rate from mobile sources was low, the collagen area fraction was reduced significantly (Fig 4H,L) compared with the previous two cases. We observed similar dynamics for case DHML (Fig 4I–L) as case D (Fig 2E–H). Despite differences in spatially averaged TGF-β concentration maxima between cases DHML and DLML (Fig 4F,J), their durations of TGF-β exposure were similar leading to very similar spatially averaged collagen concentrations (Fig 4G,K).

### Comparison between simulated and experimental studies of fibroblasts dynamics

Experimental observations of mice models reported that during MI, fibroblasts appeared around day 4, and fibroblasts and collagen deposition remained prominent from day 4 to 21 [57]. We also observed the activation and recruitment of fibroblasts around 4 days (Fig 4), and fibroblast population and collagen deposition remained prominent for a higher secretion rate of TGF-β from M2 macrophages (Fig 4A in case DHMH and Fig 4M in case DLMH). Other data have been published showing the relative abundance of fibroblasts was higher than macrophages in COVID-19 patients [69]. The relative abundance of the cell number in Delorey et al. [69] was measured using single-cell atlases. We also observed higher ratio of fibroblast to macrophage cells in the simulation cases we considered. The intercellular model for fibrosis during MI also reported similar dynamics of macrophages, fibroblasts, latent TGF-β, and collagen [44] to some of our simulation cases. However, that model focused on fibrosis in a different organ, so exact agreement was not expected.

### Longer duration of stationary TGF-β sources causes localized fibrotic pattern

We explored the impact of longer duration for both stationary and mobile sources on fibrotic outcomes: cases DA and MA (Table 3).

### Case DA: longer duration of TGF-β activation from stationary sources

Latent stores of TGF-β from damaged sites may continue to be activated for a longer duration for severe lung tissue damage. The effects of the longer duration of stationary sources (case DA) are shown in Fig 5A–D. In this case, we turned off the secretion from mobile sources (*MS* = 0) to observe the response from damaged sites alone. We increased the released TGF-β activation duration 100-fold by decreasing the secreting agent death rate (*AT*) by a factor of 100. Fig 5A shows a greater accumulation of secreting agents over time. The concentration profile of TGF-β is proportional to the number of secreting agents. The TGF-β concentration has an approximately 2-fold larger maximum value compared to all previous cases with a peak value at an earlier phase of infection (Fig 5B). The spatial profile showed localized collagen fields with high concentrations of deposited collagen (Fig 5D). However, the violin plot of the collagen area fraction in Fig 3 showed results higher than case D and intermediate collagen area fraction compared to other cases. Therefore, even with a lower collagen fraction, localized damage with high collagen deposition can lead to long-term fibrosis by the dysregulation in the duration of activation from spatially-localized latent TGF-β sources.

**Fig 5.**
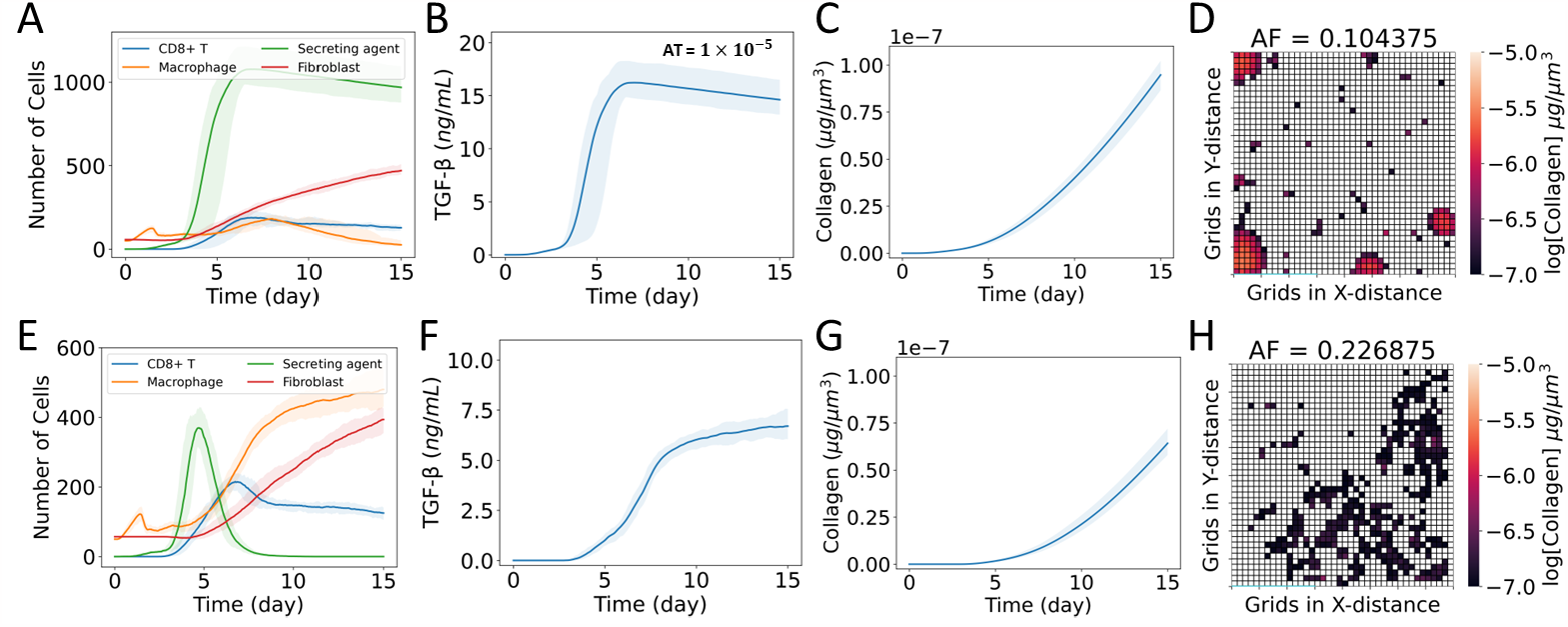
Effects of longer duration of stationary and mobile sources of TGF-β. (A–D) Case DA: longer duration of stationary damaged site sources and (E–H) case MA: longer duration of mobile macrophage sources. (A, E) Dynamics of populations of CD8+ T cells, macrophages, latent TGF-β activation sites (secreting agents), and fibroblasts; (B, F) dynamics of TGF-β concentration spatially averaged over the domain; (C, G) dynamics of collagen concentration spatially averaged over the domain; and (D, H) heat map showing collagen area fraction (*AF*) above the threshold value of 1 *×* 10^−7^ μg μm^-3^. The solid curves represent the mean of predictions, and shaded areas represent the predictions between the 5th and 95th percentile of 15 replications of the agent-based model. The cases are defined in Table 3.

We observed localization of fibroblasts around 2 days with slow creation of stationary sources (Fig S12B,I), which is disrupted around 5 days (Fig S12C,J) by the rapid creation of new stationary sources. The longer duration of stationary sources resulted in persistent localization of fibroblasts at the damaged sites from 5 days to 15 days (Fig S12C–G,I–N). The localization pattern remained the same during that period due to a slow decrease in stationary sources. The number of accumulated fibroblasts also increased in localized positions.

As a further *in silico* experiment, we changed the initial placement of virions in case DA. All cases thus far used uniform random distribution of virions. In this scenario, we considered a Gaussian distribution at the tissue center for case DA. Figs S13 and S14 showed higher fibroblast accumulation and denser collagen deposition near the center compared with the default random distribution. While only a single major fibrotic region developed (Fig S14D), the collagen concentration was the highest among all cases considered here. Thus, fibrotic patterns vary depending on the location of initial infection sites.

Our simulated results are similar to the features of focal fibrosis. Pulmonary focal fibrosis is a non-specific tissue response to injury, including infection. The feature of focal fibrosis is the sharply demarcated nodular ground-glass opacity with a maximal diameter of less than 2 cm on a thin-section computed tomography scan. The pathogenesis of focal fibrosis still needs to be explored [70]. Some studies speculated that focal fibrosis was associated with microscopic arterio-venous fistula or overexpression of platelet-derived growth factor-B [70, 71]. A multicenter observational study using 12 specimens from 11 adult COVID-19 survivors reported focal fibrosis in 50% of the patients [72]. Since we observed similar characteristic features of focal fibrosis, we hypothesized that focal fibrosis can be produced by longer activation of latent TGF-β sources.

### Case MA: no apoptosis of M2 macrophages

Sefik et al. [73] reported experimental results showing that severe SARS-CoV-2 infection increases human lung macrophage numbers at 4 days with a peak value at 14 days, and the number of macrophages remains high until 28 days. They also observed similar dynamics for the number of macrophages marked by CD206hi (a marker for M2 macrophages [74]), CD86+, and CD169+ expression. Sefik et al. [73] further reported persistent lung tissue pathology in the exudative, organizing, and fibrotic phases, with changes in affected tissue areas between 45% and 55% from uninfected tissue. We investigated whether the model could predict these findings by considering a similar scenario (case MA) by turning off the apoptosis rate of M2 macrophages (*μ*_*F*_ = 0) and the activation rate of TGF-β from damaged sites (*DS* = 0 ng/min). In Fig 5E–H, we showed the effects of longer activation of M2 macrophages. The spatially averaged collagen concentration (Fig 5G) had a similar shape, but somewhat smaller values, than that for case DA (Fig 5C), We observed a higher number of macrophages in case MA compared to other cases (Fig 5E and Fig S15). To determine the compositions of the macrophage populations reported in Figs 2 and 5, we generated Fig S16; in all four cases shown (as in the others not shown), the M2 macrophage phenotype dominates the total macrophage population as the numbers of secreting agents peak (due to death of infected epithelial cells). The simulated dynamics of M2 macrophages in case MA (Fig S16) were also similar to experimental observations of Sefik et al. [73]. In Fig 5F, the spatially averaged TGF-β concentration increased over time, remained high, and shifted its peak value compared to case DA. We also observed a higher collagen area fraction in case MA than in cases M and DA (Fig 3). The persistent presence of M2 macrophages caused a higher amount of collagen deposition, higher collagen area fraction, and localization of fibroblasts from 9 days to 15 days (Fig S15).

### Model validation for the dynamics of TGF-β, collagen area fraction, and macrophage cell population

Experimental data from the literature used in this section were extracted from graphs in the original sources (referenced in the text) using the web-based program WebPlotDigitizer [75]. Strobel et al. [76] developed a mouse model based on Adeno-associated-virus (AAV)-mediated expression of TGF-β1 to study progressive fibrosing interstitial lung diseases. The AAV-TGF-β1 model targeted TGF-β1 expression to the bronchial epithelium and alveolar type II cells, leading to persistent expression of TGF-β1 and histological features of lung fibrosis. The model characterized the dynamics of TGF-β1 protein levels in the bronchoalveolar lavage (BAL) at days 3, 7, 14, 21, and 28 after the start of the experiment. A higher level of TGF-β1 was observed on day 3 with a peak value on day 7, and the concentration of TGF-β1 remained high until 28 days (1.95–5.94 ng mL^-1^). However, they only reported the representative tissue section showing fibrotic areas at day 21. Contrary, Sefik et al. reported the percentage of affected tissue in mice models before infection and at days 2, 4, 7, 14, 28, and 35 after infection with SARS-CoV-2. However, they did not characterize the TGF-β dynamics in their experiments. Similar to the dynamics of macrophage in the experimental studies of Sefik et al. [73], Strobel et al. [76] also reported an increased number of monocytes at day 3 with a peak value at day 14, and the number of monocytes remained high until 28 days.

From our *in silico* experiments, we observed a higher concentration of TGF-β in the earlier phase of infection from the activation of latent TGF-β sources (case D) and persistent expression of TGF-β for longer activation of macrophages (case MA). Here, we considered a new case DHMA (*DS* = 2*×* 10^−9^ ng min^-1^, *MS* = 2 *×* 10^−9^ ng min^-1^, and no apoptosis of M2 macrophages) and simulated it for 28 days to validate our simulated results with experimental observations.

Our simulated result showed similar ranges and dynamics of TGF-β concentration compared to the BAL TGF-β1 protein levels in Strobel et al. [76] (Fig 6A). In addition, we observed a peak value in the TGF-β concentration around day 5. Although TGF-β concentration at day 5 was not reported, the mathematical modeling of fibrosis during MI reported peak latent TGF-β concentration at a similar time [44], and the corresponding TGF-β mRNA level in the infracted mice model reported a peak value at day 7 [77]. The dynamics of simulated collagen area fractions are also in similar ranges compared to the percentage of affected lung tissue in Sefik et al. [73] except on day 2 and day 14 (Fig 6B). In addition, we quantified collagen area fractions using the open-source platform Fiji [78] from the day 21 representative images of NaCl control and AAV-TGF-β1 treated lung tissue section in Strobel et al. [76]. We considered the area fraction from NaCl control as an uninfected condition. Contrary to the experimental studies of Strobel et al. [76], we assumed no collagen in our simulation before infection. However, we observed a similar collagen area fraction at day 21 compared to the area fraction quantified from the AAV-TGF-β1 treated tissue section in Strobel et al. [76]. To compare the simulated M2 macrophages dynamics with the experimental data, we calculated the mean at each time point *t*_*j*_ for *n* number of samples as 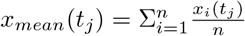 and scaled the cell count by dividing all the data with the maximum value of means (*max*(*x*_*mean*_(*t*_*j*_)). Our simulated dynamics of M2 macrophages were also similar to the population dynamics of alveolar macrophages reported in Sefik et al. [73] and monocyte dynamics reported in Strobel et al. [76] (Fig 6C).

**Fig 6.**
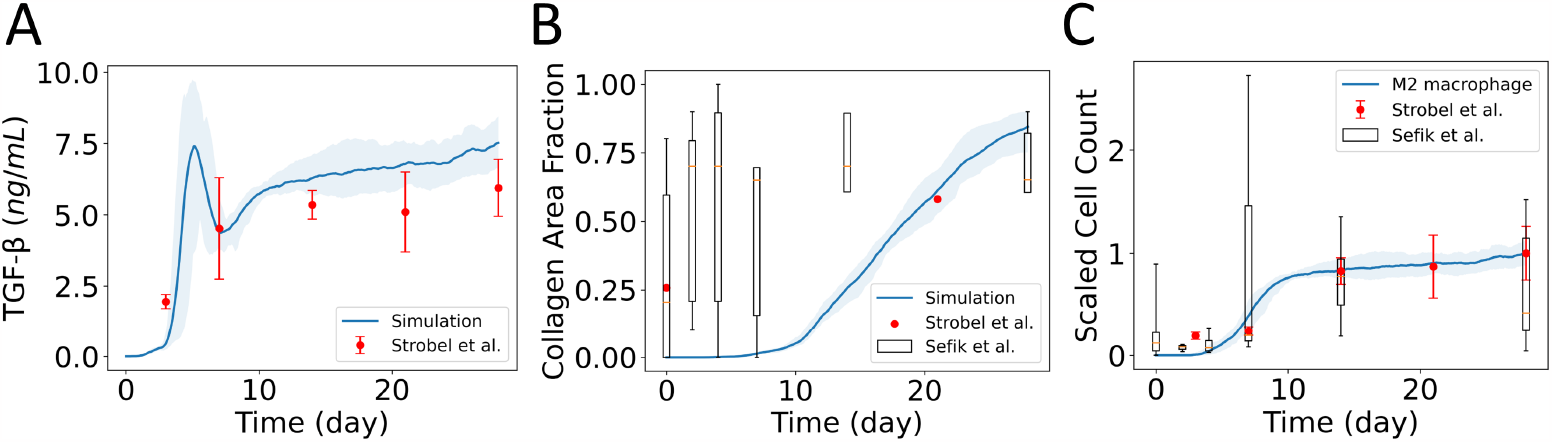
Model validation with experimental data of TGF-β, collagen, and macrophage cell population. A) Comparison between experimental data of TGF-β in Strobel et al. [76] and simulated TGF-β concentration spatially averaged over the domain. B) Comparison of simulated collagen area fractions with experimental data from Sefik et al. [73] and Strobel et al. [76]. C) Comparison of the dynamics of simulated M2 macrophage count with the population dynamics of alveolar macrophages reported in Sefik et al. [73] and monocytes reported in Strobel et al. [76]. The solid curves represent the mean of predictions, and shaded areas represent the predictions between the 5th and 95th percentile of 8 replications of the agent-based model. In the box plots from Sefik et al. [73], the whiskers go down to the smallest value (minimum) and up to the largest value (maximum), the box extends from the 25th to 75th percentiles, and the median is shown as a line inside the box.

### Partial removal of TGF-β changes fibrotic pattern

We considered the impact of TGF-β uptake by fibroblasts as they chemotax through the domain by adding an uptake term in Eq 4:

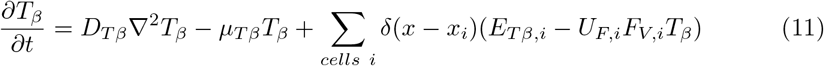

where *U*_*F,i*_ is the uptake rate of TGF-β by fibroblast cell *i* and *F*_*V,i*_ is the volume of fibroblast cell *i*. The uptake rate of TGF-β by fibroblasts was assumed equivalent to the uptake rate of debris in the overall model. Here we considered cases DAU and MAU (Table 3). Figs S17 and S18 show the population spatial distributions over time for each agent type for these two cases. Fig S19 compares the collagen area fractions for these final cases to other cases.

### Case DAU: uptake of TGF-β by fibroblasts in case DA

We considered the uptake of TGF-β in case DA and termed this scenario as case DAU. Although the population dynamics and mean collagen concentrations (Fig 7A,C) remained similar to those in case DA (Fig 5A,C), we observed variations in the spatially averaged TGF-β dynamics with a reduced peak value (Fig 7B). The uptake of TGF-β also changed the concentration gradient of TGF-β in the ECM (Eq 6), which caused delocalization and redistribution of fibroblasts in other tissue areas (Fig S17). This effectively maintained a chemotactic gradient to drive fibroblast movement. As a result, we observed a significantly higher collagen area fraction in case DAU compared to cases D and DA (Fig 7D and S19).

**Fig 7.**
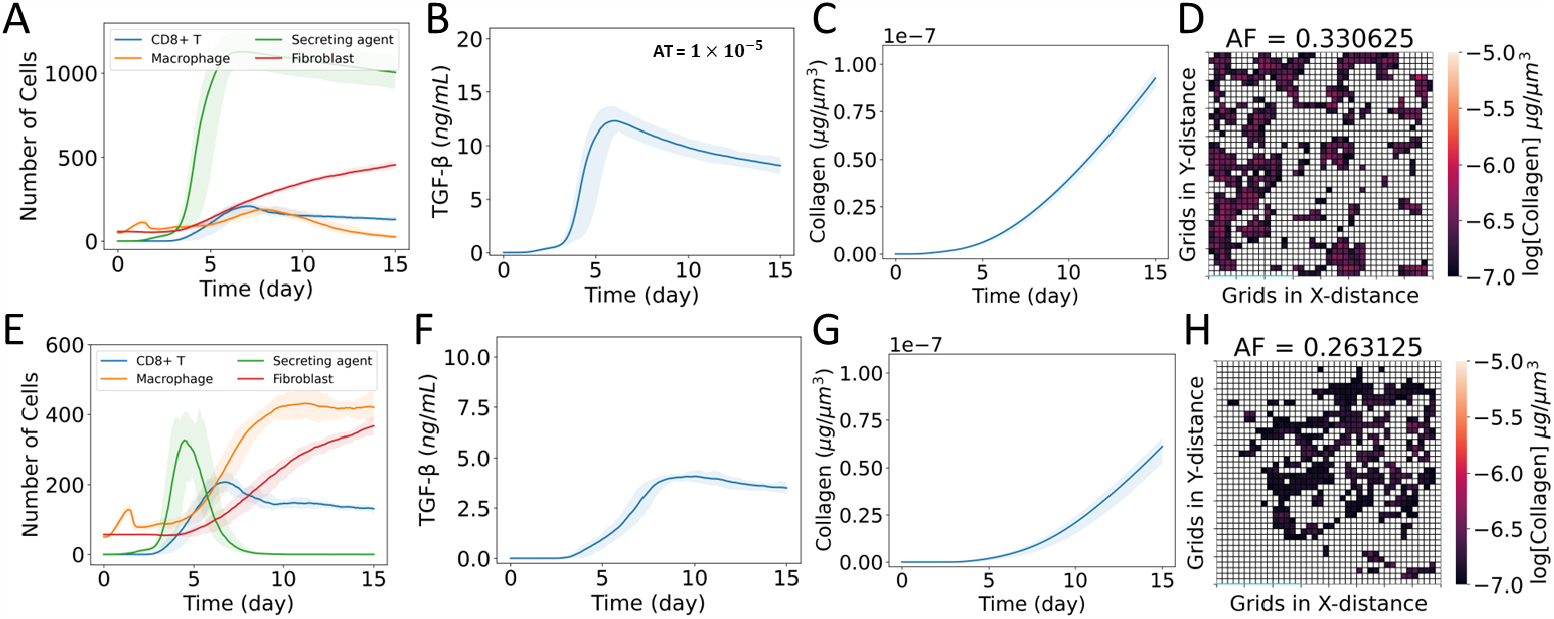
Effects of the uptake of TGF-β by fibroblasts. (A–D) Case DAU: uptake of TGF-β by fibroblasts in case DA and (E–H) case MAU: uptake of TGF-β by fibroblasts in case MA. (A, E) Dynamics of populations of CD8+ T cells, macrophages, latent TGF-β activation sites (secreting agents), and fibroblasts; (B, F) dynamics of TGF-β concentration spatially averaged over the domain; (C, G) dynamics of collagen concentration spatially averaged over the domain; and (D, H) heat map showing collagen area fraction (*AF*) above the threshold value of 1 *×* 10^−7^ μg μm^-3^. The solid curves represent the mean of predictions, and shaded areas represent the predictions between the 5th and 95th percentile of 15 replications of the agent-based model. The cases are defined in Table 3.

### Case MAU: uptake of TGF-β by fibroblasts in case MA

We also considered the uptake of TGF-β in case MA and termed that scenario case MAU. Here, we observed reductions in spatially averaged TGF-β dynamics with similar populations and spatially averaged collagen dynamics (Fig 7E–H) compared to case MA (Fig 5E–H). Although the delocalization and redistribution of fibroblasts were observed in case MAU (Fig S18), collagen area fractions remained similar to case MA (Fig S19). We also observed higher collagen area fractions in case DAU compared to case MAU. Our findings suggest that removing TGF-β from ECM when persistent latent TGF-β activation sites are present can spread fibrotic areas in tissue, which is particularly noticeable when spatially averaged TGF-β concentration is high.

### Limitations and potential future application to clinical trials

Here, we quantified the effects of stationary and mobile sources of TGF-β in the progression of lung fibrosis. Our simplified model assumed a homogeneous distribution of latent TGF-β in the tissue domain, and epithelial cell death at any location activates the latent TGF-β. However, the activation time of latent TGF-β sources is probabilistically sampled from an exponential distribution with a mean death rate of the sources to simulate the variability in the sources. We also assumed immediate activation of latent TGF-β from M2 macrophages.

The production rate of TGF-β from stationary and mobile sources was assumed constant and we did not consider the effect of cell volume on the production rate. Instead, we iteratively selected the ranges of TGF-β production rate from sources to keep extracellular TGF-β concentration within experimentally observed ranges. We are able to recreate the experimentally observed TGF-β levels in Strobel et al. [76] and dynamics without calibrating any model parameters from the experimental data. Other mathematical modeling and experimental articles also used similar ranges of TGF-β production rates [43–45].

The phenotypic shift of M1 to M2 macrophages involves several different pathways and cytokines [15]. However, our model assumed a simplified rule for the transition of M1 to M2 phenotype (Fig 1).

We started the simulation from a fixed number of inactive fibroblasts, assuming homeostatic conditions. Our model did not account for the birth and death of initial inactive fibroblasts to maintain homeostasis. We also assumed a simplified rule to consider active and inactive states of fibroblasts in the presence and absence of TGF-β, respectively. We did not include the mechanistic details of TGF-β-dependent fibroblast activation. Also, we did not account for the mechanical interactions of fibroblasts with epithelial cells and other immune cells. Our model is limited to the dynamics of fibroblasts depending on the biochemical factors secreted by other cells and tissue. We also assumed fibroblast chemotaxis along the gradients of TGF-β and did not consider the effect of other chemokines involved in the migration of fibroblasts. Our model did not consider fibroblasts’ transition to myofibroblasts, a hyperactive state of fibroblasts.

Initially, we did not consider any collagen in our tissue section. We assumed that represents the baseline condition and quantified the changes from baseline. We assumed collagen is continuously secreted from the active fibroblasts and deposited collagen is non-diffusing. We also did not account for the degradation of deposited collagen. Several studies highlighted matrix metalloproteinases (MMPs) as regulators of ECM degradation [40, 43, 44]. We assumed a threshold value of the deposited collagen to account for the effect of collagen degradation in the quantification of fibrotic areas from the simulated tissue.

The community-driven overall model was developed and cross-checked by a large coalition of math biologists, virologists, and immunologists over several rounds of open source revision [46]. We used the overall model as a baseline model to simulate the disease progression. However, the rules and parameters used in the fibrosis model are extendable to other disease models to quantify the effect of TGF-β sources in the progression of lung fibrosis.

Our results give critical insight into the current clinical trials targeting TGF-β for lung fibrosis. At present, two of the promising drugs targeting the TGF-β signaling pathway for fibrosis treatment are pirfenidone and PLN-74809, and both are in Phase II clinical trials [79]. The mechanism of pirfenidone involves inhibiting the activity of TGF-β by stopping the maturation of latent TGF-β [80, 81]. The drug PLN-74809 inhibits αvβ_6_ and αvβ_1_ integrins to prevent the activation of latent TGF-β [82]. In our simulations, we observed that (1) changing the TGF-β sources alters the fibrotic pattern and (2) removing TGF-β from the ECM in the presence of persistent activation sizes leads to increases in the collagen area fractions. If these computational findings can be validated, the results have important implications for ongoing clinical trials: improving the trial outcomes requires a more thorough consideration of the spatial behavior of fibroblasts and immune cells and the rate and duration of TGF-β activation. Partial removal of TGF-β from ECM or inhibition of TGF-β activation from sources may change the chemotactic direction of fibroblasts and lead to fibrotic areas with higher collagen area fraction. It is essential to consider the spatial behavior and activity of sources before targeting them for inhibition of TGF-β as fibrosis is a spatial phenomenon.

## Conclusions

The multiscale tissue simulator (overall model and fibrosis model) described here gives insight into the dynamics and impact of TGF-β sources in fibroblast-mediated collagen deposition at damaged sites of SARS-CoV-2 infected tissue in COVID-19. We applied this model to investigate the spatial behaviors of sources of TGF-β, the activation rate of TGF-β, and the activation duration of TGF-β sources in the process of lung fibrosis. Fibrosis is a spatial event; thus, it is essential to consider the spatial properties of the sources of the key biomarkers observed during the process. Our results showed localized collagen deposition from stationary sources of TGF-β and dispersed collagen deposition from mobile sources of TGF-β. Our results indicated that M2 macrophages and their persistent presence were mainly responsible for higher collagen area fractions. The computational observations were supported by independent experimental and clinical observations. We predicted a longer duration of stationary TGF-β sources could lead to fibrosis even with a lower collagen area fraction. Also, partial removal of TGF-β from ECM changed the chemotactic gradient of fibroblasts, resulting in a higher collagen area fraction. In the future, the accuracy of the predictions could further be improved by considering the premorbidity of patient groups (e.g., hypertension), different phenotypes of fibroblasts (e.g., myofibroblasts), negative regulators for collagen degradation (e.g., MMP9), and collagen feedback to epithelial cells for regrowth and wound healing and to immune cells for transport properties through the evolving ECM. The quantification of the collagen deposition dynamics depending on the activity of sources may benefit treatment strategies controlling TGF-β and help characterize the evolution of fibrosis in COVID-19 survivors.

## Supporting information

### Supplementary File

The Supplementary File contains supporting information and figures organized into the following sections: Agent-based model decisions and workflow, TGF-β-dependent functions for fibroblasts (Fig S1); fibroblast collagen production rate *k*_*F C*_; effects of TGF-β activation/secretion rate from each source (Fig S2); dynamics of virions, neutrophils, dendritic cells, and CD4 cells (Fig S3); effects of turning on/off TGF-β activation and secretion rates from stationary and mobile sources (Figs S4–S7); effects of varying the TGF-β activation rate from stationary damaged sites and the secretion rate from mobile macrophages at different levels in combination (Figs S8–S11); effects of increasing the duration of the sources (Figs S12–S16); and effects of uptake of TGF-β (Figs S17–S19).

## Acknowledgments

This work was supported by the National Institutes of Health grant R35GM133763 (to A.N.F.V.), the University at Buffalo, and the Center for Computational Research at the University at Buffalo. We thank the scientific community for fibrosis model feedback, particularly Amber M. Smith, Thomas Hillen, Yafei Wang, and members of the Ford Versypt Lab.

