## Supplementary File for "An agent-based modeling approach for lung fibrosis in response to COVID-19"

### Agent-based model decisions and workflow

#### Overall model

The overall model has 5 cell types and 76 rules. Here, we list all the cell types and biological hypotheses (agent-based model rules) from the overall model manuscript [1].

##### Epithelial cells

1. Live epithelial cells undergo apoptosis after sufficient cumulative contact time with adhered CD8+ T cells
2. Dead epithelial cells produce debris
3. Virus adheres to unbound external ACE2 receptor to become external (virus)-bound ACE2 receptor
4. Bound external ACE2 receptor is internalized (endocytosed) to become internal bound ACE2 receptor
5. Internalized bound ACE2 receptor releases its virion and becomes unbound internalized receptor. The released virus is available for use by the viral lifecycle model
6. Internalized unbound ACE2 receptor is returned to the cell surface to become external unbound receptor
7. Each receptor can bind to at most one virus particle
8. Internalized virus is uncoated
9. Uncoated virus (viral contents) lead to release of functioning RNA
10. RNA creates viral protein at a constant rate unless it degrades
11. Viral RNA is replicated at a rate that saturates with the amount of viral RNA
12. Viral RNA undergoes constitutive (first order) degradation
13. Viral protein is transformed to an assembled virus state

14. Assembled virus is released by the cell (exocytosis)
15. After infection, cells secrete chemokine
16. As a proxy for viral disruption of the cell, the probability of cell death increases with the total number of assembled virions
17. Apoptosed cells lyse and release some or all their contents
18. Once viral RNA exceeds a particular threshold, the cell enters the pyroptosis cascade
19. Once pyropotosis begins, the intracellular cascade is modeled by a system of ODEs monitoring cytokine production and cell volume swelling
20. Cell secretion rate for pro-inflammatory increases to include secretion rate of IL-18
21. Cell secretes IL-1$\beta$ which causes a bystander effect initiating pyroptosis in neighboring cells
22. Cell lyses (dying and releasing its contents) once its volume has exceeded $1.5\times$ the homeostatic volume
23. Infected epithelial cells secrete pro-inflammatory cytokine
24. Antigen presentation in infected cells is a function of intracellular viral protein

**Macrophages**

1. Resident (unactivated) and newly recruited macrophages move along debris gradients
2. Macrophages phagocytose dead cells. Time taken for material phagocytosis is proportional to the size of the debris
3. Macrophages break down phagocytosed materials
4. After phagocytosing dead cells, macrophages activate and secrete pro-inflammatory cytokines
5. Activated macrophages can decrease migration speed
6. Activated macrophages have a higher apoptosis rate
7. Activated macrophages migrate along chemokine and debris gradients
8. Macrophages are recruited into tissue by pro-inflammatory cytokines
9. Macrophages can die and become dead cells only if they are in an exhausted state
10. Macrophages become exhausted (stop phagocytosing) if internalized debris is above a threshold
11. CD4+ T cell contact induces activated macrophage phagocytosis of live infected cells

**Neutrophils**

1. Neutrophils are recruited into the tissue by pro-inflammatory cytokines
2. Neutrophils die naturally and become dead cells
3. Neutrophils migrate locally in the tissue along chemokine and debris gradients
4. Neutrophils phagocytose dead cells and activate

5. Neutrophils break down phagocytosed materials
6. Activated neutrophils reduce migration speed
7. Neutrophils uptake virus
8. Neutrophils secrete ROS upon phagocytosis

**Dendritic cells (DCs)**

1. Resident DCs exist in the tissue
2. DCs are activated by infected cells and/or virus
3. Portion of activated DCs leave the tissue to travel to the lymph node
4. DCs chemotaxis up chemokine gradient
5. Activated DCs present antigen to CD8+ T cells increasing their proliferation rate and killing efficacy (doubled proliferation rate and attachment rate)
6. Activated DCs also regulate the CD8+ T cell levels in within a threshold by enhancing CD8+ T cell clearance

**CD8+ T cells**

1. CD8+ T cells are recruited into the tissue by pro-inflammatory cytokines
2. CD8+ T cells apoptose naturally and become dead cells
3. CD8+ T cells move locally in the tissue along chemokine gradients
4. CD8+ T cells adhere to infected cells. Cumulated contact time with adhered CD8+ T cells can induce apoptosis
5. Activated DCs present antigen to CD8+ T cells, which increases the CD8+ T cell proliferation rate
6. Activated DCs also regulate the CD8+ T cell levels in within a threshold by enhancing CD8+ T cell clearance
7. CD8+ T cells have a max generation counter and will not proliferate after the set generation

**CD4+ T cells**

1. CD4+ T cells are recruited into the tissue by the lymph node
2. CD4+ T cells apoptose naturally and become dead cells
3. CD4+ T cells move locally in the tissue along chemokine gradients
4. CD4+ T cells are activated in the lymph node by three signals: antigenic presentation by the DCs, direct activation by cytokines secreted by DCs, and direct activation by cytokines secreted by CD4+ T cells
5. CD4+ T cells are suppressed directly by cytokines secreted by CD4+ T cells
6. CD4+ T cells have a max generation counter and will not proliferate after the set generation

**Tissue microenvironment**

1. Virus diffuses in the microenvironment
2. Virus adhesion to a cell stops its diffusion (acts as an uptake term)
3. Pro-inflammatory cytokine diffuses in the microenvironment
4. Pro-inflammatory cytokine is taken up by recruited immune cells

5. Pro-inflammatory cytokine is eliminated or cleared
6. Chemokine diffuses in the microenvironment
7. Chemokine is taken up by immune cells during chemotaxis
8. Chemokine is eliminated or cleared
9. Debris diffuses in the microenvironment
10. Debris is taken up by macrophages and neutrophils during chemotaxis
11. Debris is eliminated or cleared
12. Immunoglobulin (Ig) diffuses in the microenvironment
13. Virions and Ig react in tissue removing both components
14. Reactive oxidative species (ROS) diffuses in the microenvironment

#### Fibrosis model

We extended the overall model to include the mechanisms of fibrosis. The details of the cell types and biological hypotheses are described in detail in the manuscript. Here, we summarize the cell types and biological hypotheses (agent-based model rules) for the fibrosis model.

##### M2 Macrophages

1. CD8+ T cell contact stops activated macrophage secretion of pro-inflammatory cytokine and switches to M2 phase
2. M2 macrophages secrete TGF-β

##### Secreting agent

1. Secreting agents are created at the site that a CD8+ T cell kills an epithelial cell to mimic latent TGF-β activation embedded in the tissue
2. TGF-β cytokine is eliminated or cleared
3. Secreting agents secrete TGF-β for a set amount of time

##### Fibroblasts

1. Resident inactive fibroblasts exist in the tissue
2. Resident inactive fibroblasts do not apoptose to maintain homeostasis
3. TGF-β activates the resident inactive fibroblasts and absence of TGF-β inactivates fibroblasts
4. Active fibroblasts are recruited into the tissue by TGF-β
5. Active fibroblasts and inactive fibroblasts from the active state apoptose naturally and become dead cells
6. Fibroblasts move locally in the tissue along up gradients of TGF-β
7. Fibroblast cells deposit collagen continuously

##### Tissue microenvironment

1. TGF-β diffuses in the microenvironment
2. TGF-β is degraded or removed
3. Collagen does not diffuse and degrade

#### TGF-β-dependent functions for fibroblasts

The function for TGF-β-dependent recruitment of fibroblasts from [2] was fit to experimental data of [3], represented by solid red circles, and is shown in Fig S1A. This fitted curve is used as $F_g(T_\beta)$ for recruitment of new fibroblasts in Eq 5 of the fibrosis model. We assumed constant response (green line) when TGF-β concentration exceeds the threshold value (10 ng/mL, marked by the vertical black line in Fig S1A) because of the polynomial nature of the recruitment signal.

Fig S1B shows the function used for TGF-β-dependent collagen deposition rate from fibroblasts, $F_c(T_\beta)$. The experimental data from [4,5], represented by solid red circles, were normalized and fitted with a Michaelis-Menten kinetic function represented by a blue curve to estimate $V_{T\beta}$ and $k_{T\beta}$ for $F_c(T_\beta)$ in Eq 7 of the fibrosis model.

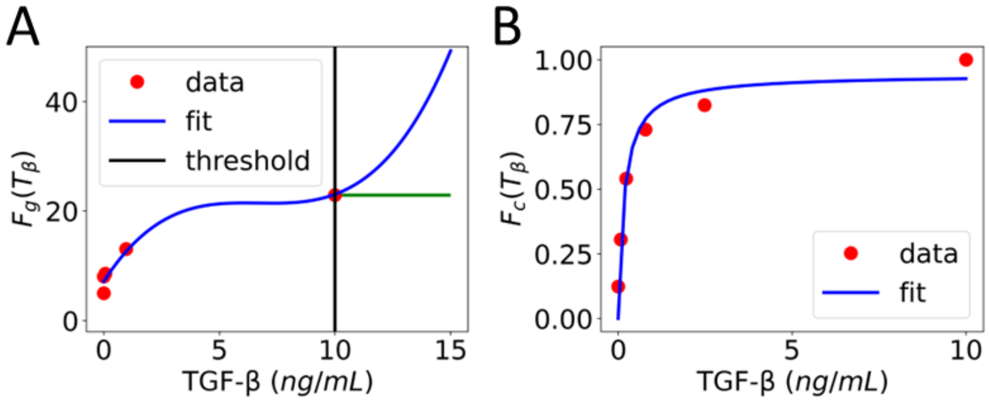


**Fig S1.** TGF-β concentration dependency for functions used in fibrosis model to describe (A) recruitment of fibroblasts (Eq 5) and (B) collagen deposition from fibroblasts (Eq 7). In A) the green curve is used as the $F_g(T_\beta)$ constant value for $T_\beta > 10$.

#### Fibroblast collagen production rate $k_{FC}$

We estimated $k_{FC}$ from Fig 3 of Hao et al. [6]. We extracted data between day 4 and day 9 of the fibroblast population and the ECM concentration because we observed fibroblast activation and recruitment in our model during that period. Also, in Hao et al. [6], the population dynamics and ECM deposition remained linear during that period. We assumed fibroblast cell weight was $2 \times 10^{-9}$ g to calculate the number of fibroblasts. We used Eq S1 to estimate fibroblast collagen production rate ($k_{FC}$):

$$k_{FC} = \frac{\text{slope of ECM}}{\#\text{ of fibroblasts} * \text{time interval}} \tag{S1}$$

#### Effects of TGF-β activation/secretion rate from each source

The fibrosis model considers two different sources that can activate TGF-β. The first source activates TGF-β from latent stores in the tissue ECM at the stationary damaged sites, and the rate of this activation is denoted $DS$ for the damaged site. The second source is from the secretion and activation of TGF-β from mobile M2 macrophages, and the rate of this secretion is denoted $MS$ for macrophage secretion. In Fig S2 the impacts of a range of values for $DS$ and $MS$ are considered from either source individually, but not in combination. $DS$ or $MS$ values of 0 represent when the respective source is effectively turned off in the simulations. Values of $DS$ and $MS$ deemed acceptable were those that kept TGF-β concentrations below 10 ng/mL while also still have some visually noticeable TGF-β concentrations within the 15 day time period.

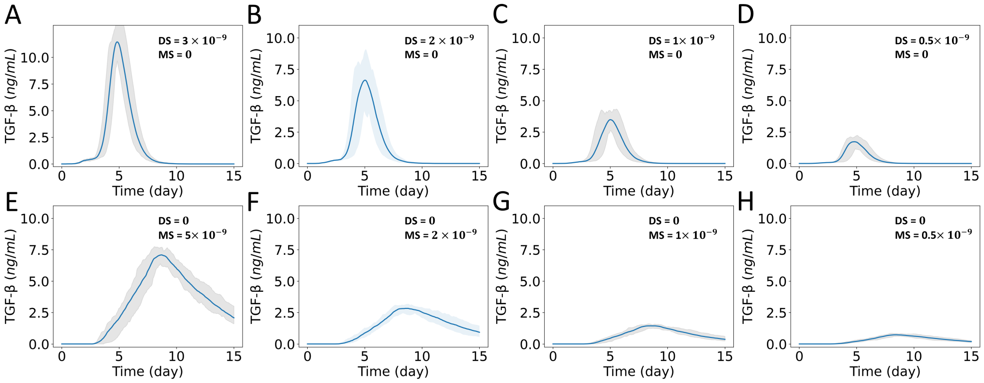


**Fig S2.** Changes in the dynamics of TGF-β concentration with variations in the activation rate of latent TGF-β from damaged sites ($DS$) and secretion rate from mobile macrophages ($MS$). Dynamics of TGF-β spatially averaged over the domain for $DS$ values of (A) $3 \times 10^{-9}$, (B) $2 \times 10^{-9}$, (C) $1 \times 10^{-9}$, and (D) $0.5 \times 10^{-9}$ ng/min while $MS = 0$, and for $MS$ values of (E) $5 \times 10^{-9}$, (F) $2 \times 10^{-9}$, (G) $1 \times 10^{-9}$, and (H) $0.5 \times 10^{-9}$ ng/min while $DS = 0$. The solid curves represent the mean of predictions, and shaded areas represent the predictions between the 5th and 95th percentile of 15 replications of the agent-based model.

#### Dynamics of virions, neutrophils, DCs, and CD4 cells

The overall model produces predictions for the viral virions and immune cells: neutrophils, dendritic cells (DC), and CD4+ T Cells. The dynamics of these species are inputs to the fibrosis model. The fibrosis model output does not feedback to interact with any of these species directly. Details are available in Getz et al. [1].

#### Effects of turning on/off TGF-β activation and secretion rates from stationary and mobile sources

In this section, the effects of equal values for $DS$ and $MS$ are considered in combination for case DM, for just stationary damaged sites in case D, and for just mobile macrophages in case M. The values for $DS$ and $MS$ in each case are summarized in Table 3.

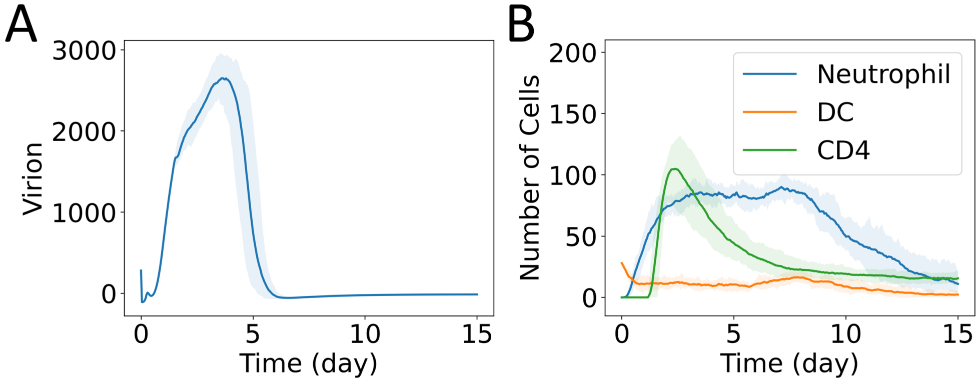


**Fig S3.** Dynamics of populations from the overall model. (A) Virions and (B) immune cells: neutrophils, dendritic cells (DC), and CD4+ T cells. The solid curves represent the mean of predictions, and shaded areas represent the predictions between the 5th and 95th percentile of 15 replications of the agent-based model.

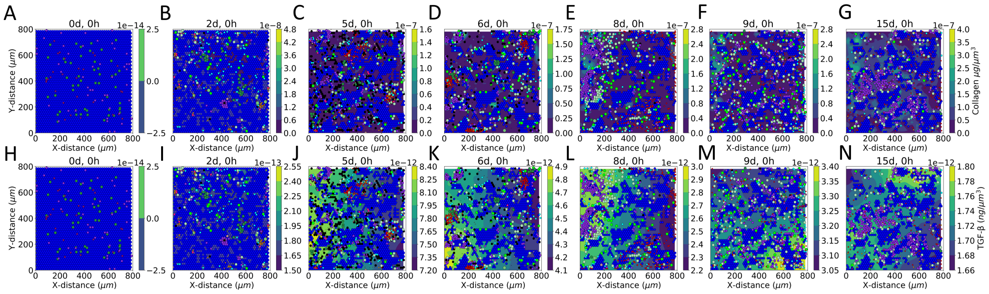

**Fig S4.** Case DM: equal TGF-β activation rate from stationary damaged sites and secretion rate from mobile macrophages. Simulated dynamics of cell population with (A–G) collagen and (H–N) TGF-β concentration fields, shown behind the cells (circles) and corresponding to the color bars. Representative model results (one replicate) at days (A, H) 0, (B, I) 2, (C, J) 5, (D, K) 6, (E, L) 8, (F, M) 9, and (G, N) 15. Epithelial cells are blue, macrophages are green, CD8+ T cells are red, secreting agents are black, and fibroblasts are purple. The color codes for other immune cells are the same as in Getz et al. [1]. The cases are defined in Table 3.

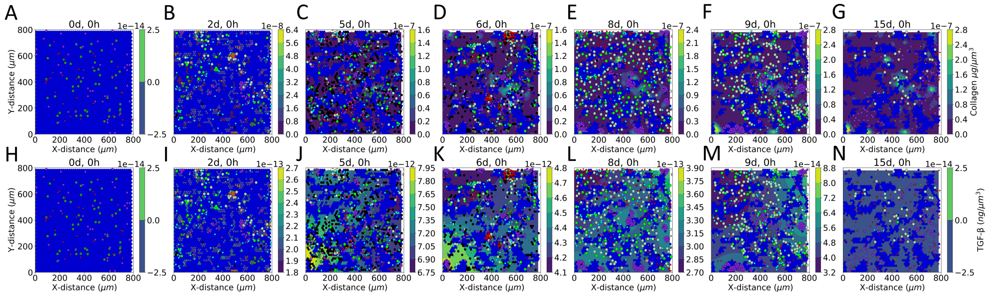


**Fig S5.** Case D: TGF-β activation from stationary damaged sites only, and secretion from mobile macrophages is turned off. Simulated dynamics of cell population with (A–G) collagen and (H–N) TGF-β concentration fields, shown behind the cells (circles) and corresponding to the color bars. Representative model results (one replicate) at days (A, H) 0, (B, I) 2, (C, J) 5, (D, K) 6, (E, L) 8, (F, M) 9, and (G, N) 15. Epithelial cells are blue, macrophages are green, CD8+ T cells are red, secreting agents are black, and fibroblasts are purple. The color codes for other immune cells are the same as in Getz et al. [1]. The cases are defined in Table 3.

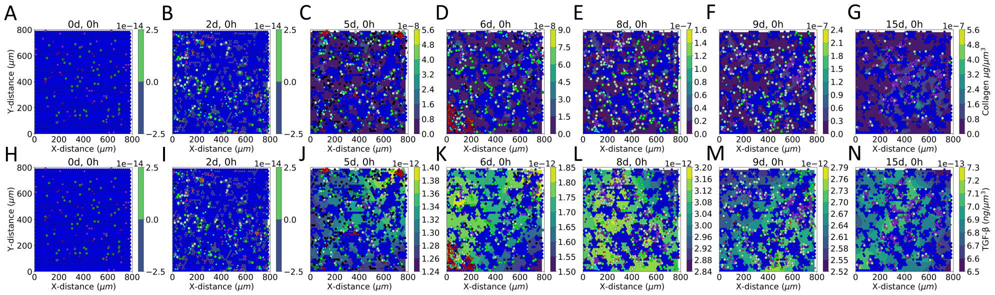


**Fig S6.** Case M: TGF-β secretion from mobile macrophages only, and activation rate from stationary damaged sites is turned off. Simulated dynamics of cell population with (A–G) collagen and (H–N) TGF-β concentration fields, shown behind the cells (circles) and corresponding to the color bars. Representative model results (one replicate) at days (A, H) 0, (B, I) 2, (C, J) 5, (D, K) 6, (E, L) 8, (F, M) 9, and (G, N) 15. Epithelial cells are blue, macrophages are green, CD8+ T cells are red, secreting agents are black, and fibroblasts are purple. The color codes for other immune cells are the same as in Getz et al. [1]. The cases are defined in Table 3.

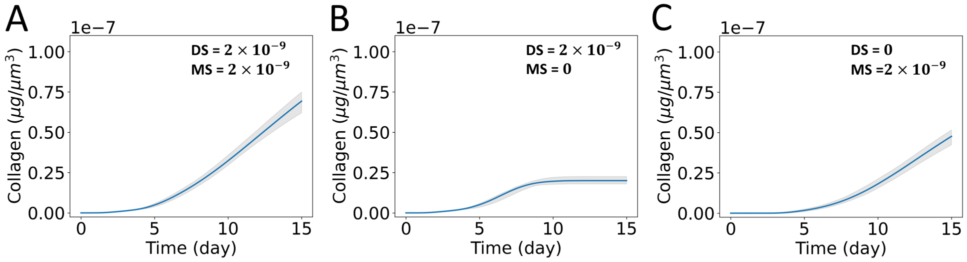


**Fig S7.** Effects of sources on collagen deposition. Dynamics of collagen concentration spatially averaged over the domain varying with the sources of TGF-β from (A) case DM, (B) case D, and (C) case M. The solid curves represent the mean of predictions, and shaded areas represent the predictions between the 5th and 95th percentile of 15 replications of the agent-based model. The cases are defined in Table 3.

### Effects of varying the TGF-β activation rate from stationary damaged sites and the secretion rate from mobile macrophages at different levels in combination

In this section the effects of combinations of values for $DS$ and $MS$ are considered for cases DHMH, DLML, DHML, and DLMH, where H and L denote higher and lower values, respectively, of the TGF-β activation rate from stationary damaged sites, denoted D, and of the TGF-β secretion rate from mobile macrophages, denoted M. The values for $DS$ and $MS$ in each case are summarized in Table 3.

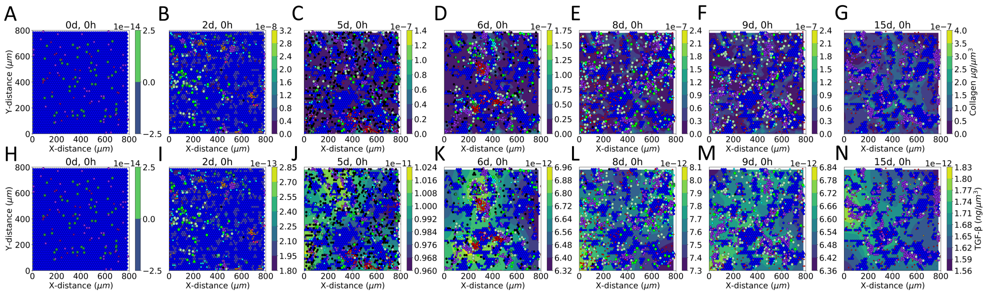


**Fig S8.** Case DHMH: high TGF-β activation rate from stationary damaged sites and high TGF-β secretion rate from mobile macrophages. Simulated dynamics of cell population with (A–G) collagen and (H–N) TGF-β concentration fields, shown behind the cells (circles) and corresponding to the color bars. Representative model results (one replicate) at days (A, H) 0, (B, I) 2, (C, J) 5, (D, K) 6, (E, L) 8, (F, M) 9, and (G, N) 15. Epithelial cells are blue, macrophages are green, CD8+ T cells are red, secreting agents are black, and fibroblasts are purple. The color codes for other immune cells are the same as in Getz et al. [1]. The cases are defined in Table 3.

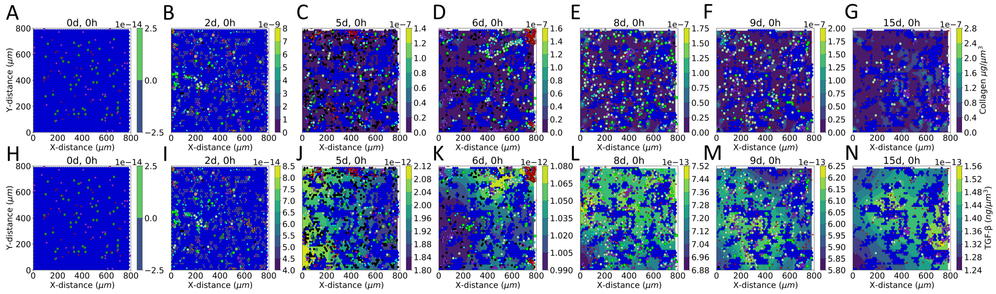


**Fig S9.** Case DLML: low TGF-β activation rate from stationary damaged sites and low TGF-β secretion rate from mobile macrophages. Simulated dynamics of cell population with (A–G) collagen and (H–N) TGF-β concentration fields, shown behind the cells (circles) and corresponding to the color bars. Representative model results (one replicate) at days (A, H) 0, (B, I) 2, (C, J) 5, (D, K) 6, (E, L) 8, (F, M) 9, and (G, N) 15. Epithelial cells are blue, macrophages are green, CD8+ T cells are red, secreting agents are black, and fibroblasts are purple. The color codes for other immune cells are the same as in Getz et al. [1]. The cases are defined in Table 3.

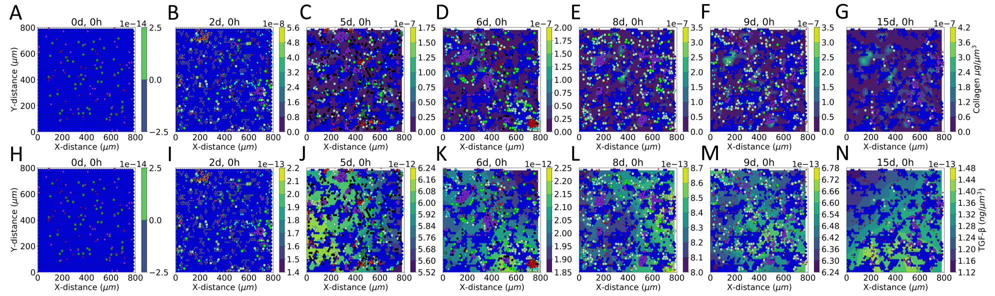


**Fig S10.** Case DHML: high TGF-β activation rate from stationary damaged sites and low TGF-β secretion rate from mobile macrophages. Simulated dynamics of cell population with (A–G) collagen and (H–N) TGF-β concentration fields, shown behind the cells (circles) and corresponding to the color bars. Representative model results (one replicate) at days (A, H) 0, (B, I) 2, (C, J) 5, (D, K) 6, (E, L) 8, (F, M) 9, and (G, N) 15. Epithelial cells are blue, macrophages are green, CD8+ T cells are red, secreting agents are black, and fibroblasts are purple. The color codes for other immune cells are the same as in Getz et al. [1]. The cases are defined in Table 3.

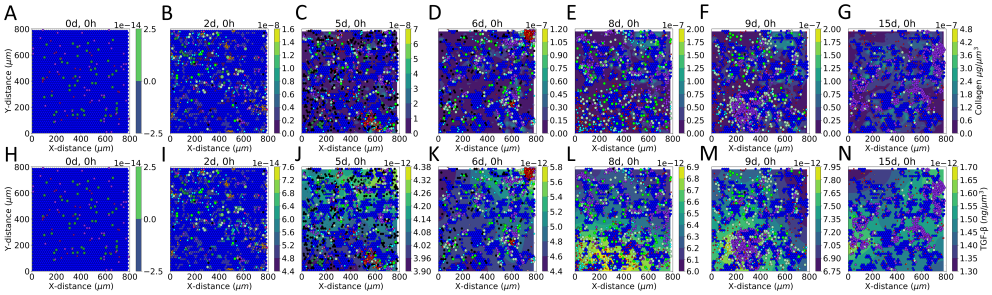

**Fig S11.** Case DLMH: low TGF-β activation rate from stationary damaged site and high TGF-β secretion rate from mobile macrophages. Simulated dynamics of cell population with (A–G) collagen and (H–N) TGF-β concentration fields, shown behind the cells (circles) and corresponding to the color bars. Representative model results (one replicate) at days (A, H) 0, (B, I) 2, (C, J) 5, (D, K) 6, (E, L) 8, (F, M) 9, and (G, N) 15. Epithelial cells are blue, macrophages are green, CD8+ T cells are red, secreting agents are black, and fibroblasts are purple. The color codes for other immune cells are the same as in Getz et al. [1]. The cases are defined in Table 3.

### Effects of increasing the duration of the sources

In this section the duration for which sources produce TGF-β is varied, and the effects are plotted. Specifically, in case DA the duration for producing TGF-β from stationary damaged sites is extended by reducing the death rate of secreting agents. In the case MA, the duration for producing TGF-β from M2 macrophages is extended by setting the apoptosis rate of M2 macrophages to 0, i.e., they do not die during the simulation time. The values for parameters in each case are summarized in Table 3.

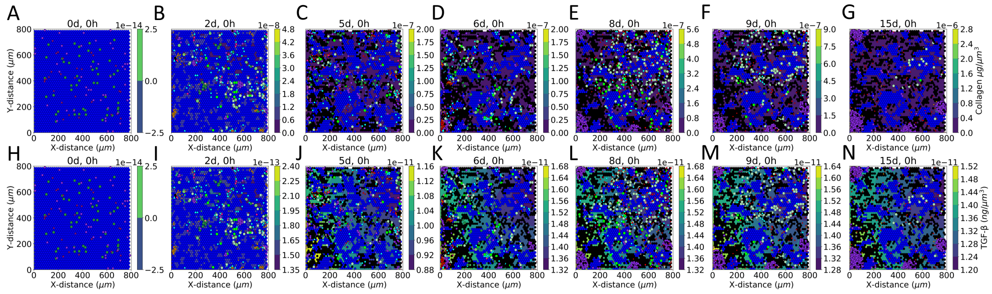

**Fig S12.** Case DA: extension of case D with longer duration of TGF-β activation from stationary damaged sites. Simulated dynamics of cell population with (A–G) collagen and (H–N) TGF-β concentration fields, shown behind the cells (circles) and corresponding to the color bars. Representative model results (one replicate) at days (A, H) 0, (B, I) 2, (C, J) 5, (D, K) 6, (E, L) 8, (F, M) 9, and (G, N) 15. Epithelial cells are blue, macrophages are green, CD8+ T cells are red, secreting agents are black, and fibroblasts are purple. The color codes for other immune cells are the same as in Getz et al. [1]. The cases are defined in Table 3.

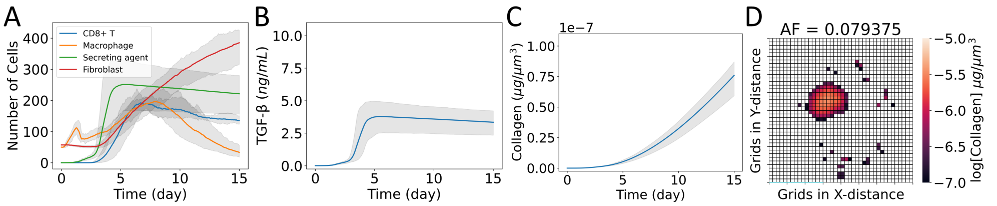


**Fig S13.** Case DA with Gaussian distribution for initial placement of virions. (A) Cell populations; (B) TGF-β concentration spatially averaged over the domain; (C) collagen concentration spatially averaged over the domain; and (D) heat map showing collagen area fraction ($AF$) above the threshold value of $1 \times 10^{-7}$ μg μm$^{-3}$. The solid curves represent the mean of predictions, and shaded areas represent the predictions between the 5th and 95th percentile of 15 replications of the agent-based model. The cases are defined in Table 3.

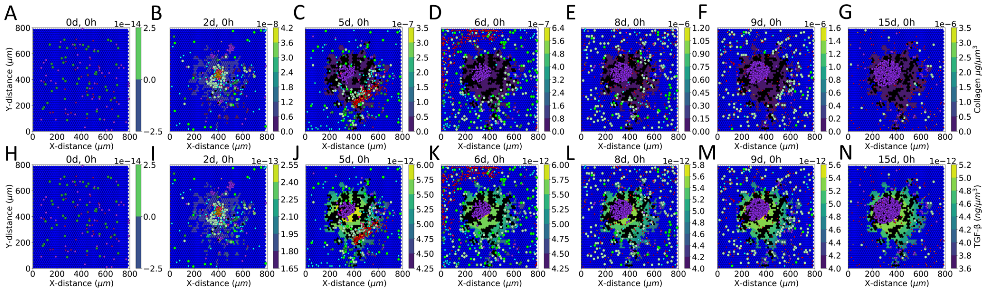


**Fig S14.** Case DA with Gaussian distribution for initial placement of virions. Simulated dynamics of cell population with (A–G) collagen and (H–N) TGF-β concentration fields, shown behind the cells (circles) and corresponding to the color bars. Representative model results (one replicate) at days (A, H) 0, (B, I) 2, (C, J) 5, (D, K) 6, (E, L) 8, (F, M) 9, and (G, N) 15. Epithelial cells are blue, macrophages are green, CD8+ T cells are red, secreting agents are black, and fibroblasts are purple. The color codes for other immune cells are the same as in Getz et al. [1]. The cases are defined in Table 3.

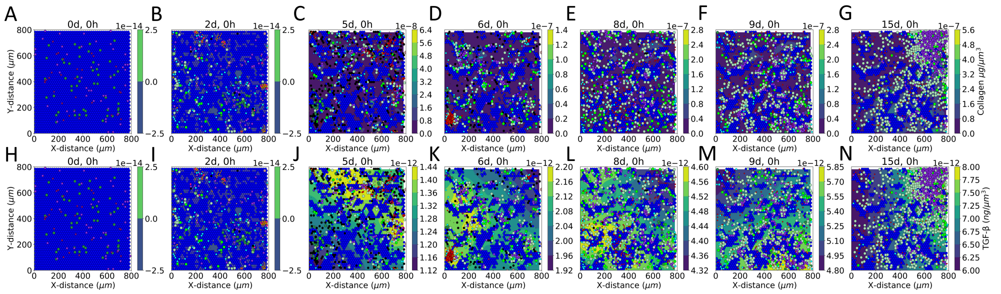


**Fig S15.** Case MA: extension of case M with longer duration of TGF-β secretion from mobile macrophages. Simulated dynamics of cell population with (A–G) collagen and (H–N) TGF-β concentration fields, shown behind the cells (circles) and corresponding to the color bars. Representative model results (one replicate) at days (A, H) 0, (B, I) 2, (C, J) 5, (D, K) 6, (E, L) 8, (F, M) 9, and (G, N) 15. Epithelial cells are blue, macrophages are green, CD8+ T cells are red, secreting agents are black, and fibroblasts are purple. The color codes for other immune cells are the same as in Getz et al. [1]. The cases are defined in Table 3.

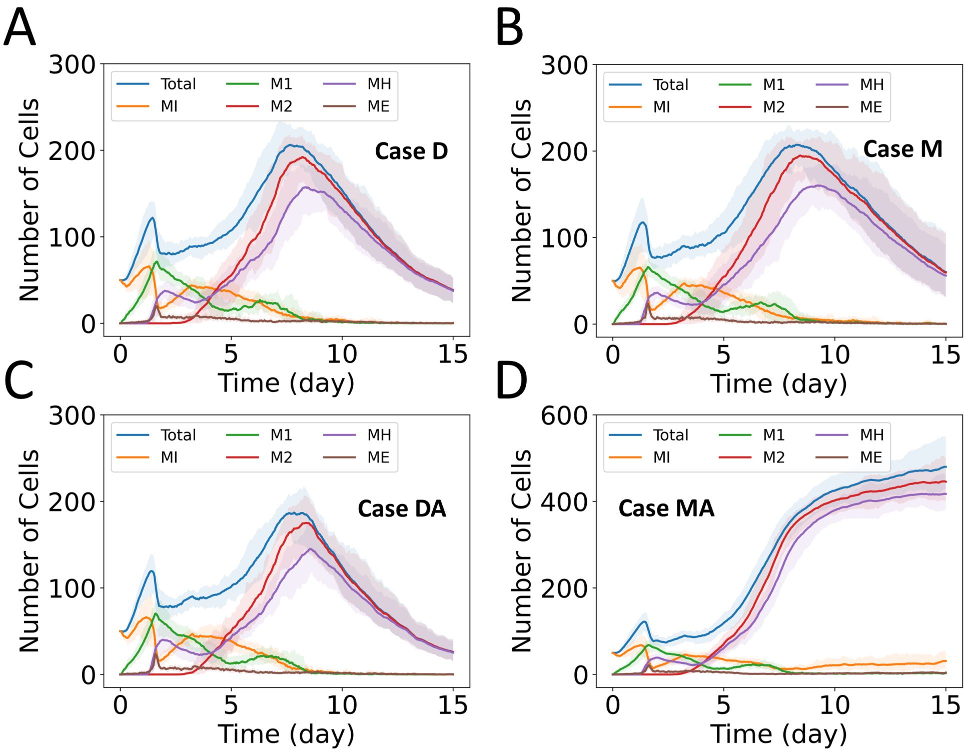


**Fig S16.** Dynamics of populations of macrophages in different states. The overall model has five states: inactivated state (MI), M1 phenotype (M1), M2 phenotype (M2), hyperactive state (MH), and exhausted state (ME). Both M1 and M2 phenotypes can also exist in MH and ME states. Details are available in Getz et al. [1]. (A) Case D, (B) case M, (C) case DA, and (D) case MA. The solid curves represent the mean of predictions, and shaded areas represent the predictions between the 5th and 95th percentile of 15 replications of the agent-based model. The cases are defined in Table 3.

### Effects of uptake of TGF-β

In this section, the effects of allowing fibroblasts to uptake TGF-β are plotted. The values for parameters in each case are summarized in Table 3.

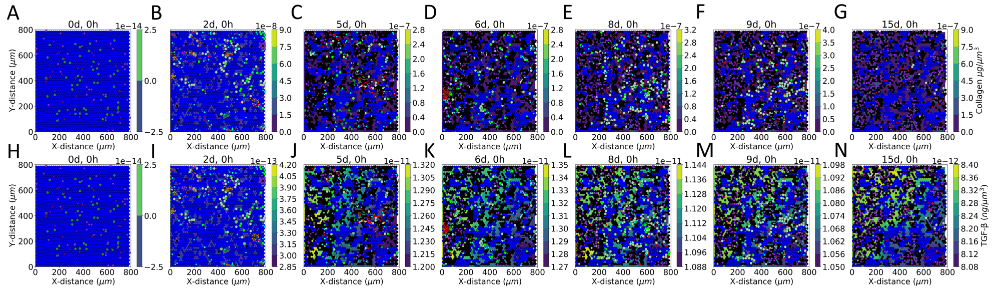


**Fig S17.** Case DAU: extension of case DA with the uptake of TGF-β by fibroblasts. Simulated dynamics of cell population with (A–G) collagen and (H–N) TGF-β concentration fields, shown behind the cells (circles) and corresponding to the color bars. Representative model results (one replicate) at days (A, H) 0, (B, I) 2, (C, J) 5, (D, K) 6, (E, L) 8, (F, M) 9, and (G, N) 15. Epithelial cells are blue, macrophages are green, CD8+ T cells are red, secreting agents are black, and fibroblasts are purple. The color codes for other immune cells are the same as in Getz et al. [1]. The cases are defined in Table 3.

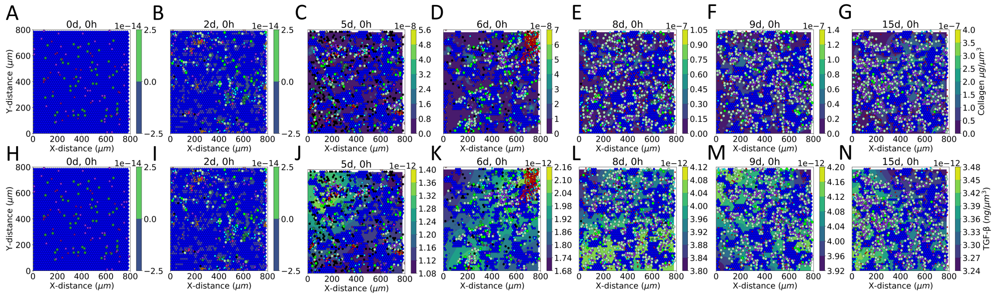


**Fig S18.** Case MAU: extension of case MA with the uptake of TGF-β by fibroblasts. Simulated dynamics of cell population with (A–G) collagen and (H–N) TGF-β concentration fields, shown behind the cells (circles) and corresponding to the color bars. Representative model results (one replicate) at days (A, H) 0, (B, I) 2, (C, J) 5, (D, K) 6, (E, L) 8, (F, M) 9, and (G, N) 15. Epithelial cells are blue, macrophages are green, CD8+ T cells are red, secreting agents are black, and fibroblasts are purple. The color codes for other immune cells are the same as in Getz et al. [1]. The cases are defined in Table 3.

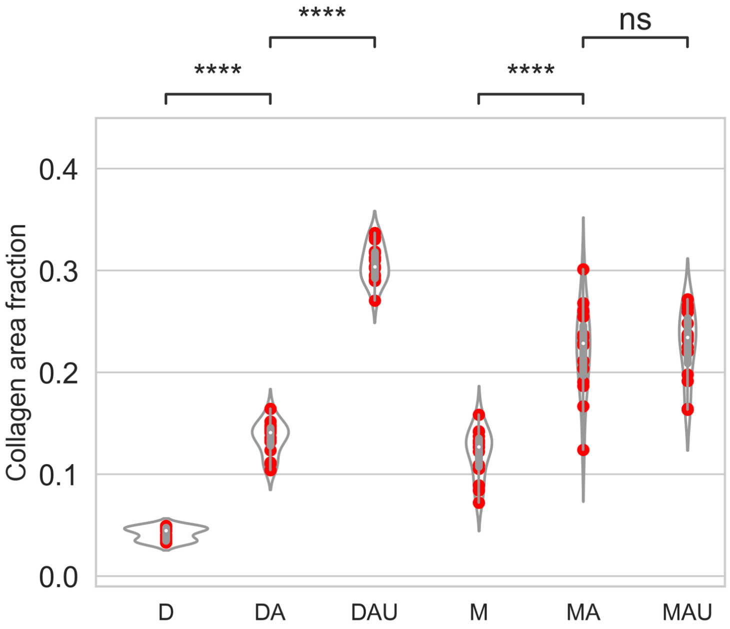


**Fig S19.** Comparison of collagen area fraction for cases D, DA, DAU, M, MA, and MAU. Violin plots represent distribution, red solid circles represent data for 15 replications of each of the cases, and box plots show medians and quartiles. The cases are defined in Table 3. $p$ values are represented by: not significant (ns) $p > 0.05$; $*p \leq 0.05$; $**p \leq 0.01$; $***p \leq 0.001$; $****p \leq 0.0001$.
